# Striatin plays a major role in angiotensin II-induced cardiomyocyte and cardiac hypertrophy in mice *in vivo*

**DOI:** 10.1101/2023.10.21.563397

**Authors:** JJ Cull, STE Cooper, HO Alharbi, SP Chothani, OJL Rackham, DN Meijles, PR Dash, Risto Kerkelä, N Ruparelia, PH Sugden, A Clerk

**Author notes:** Corresponding author: Angela Clerk, School of Biological Sciences, University of Reading, Reading, Berkshire, UK. ORCID ID: 0000-0002-5658-0708. Institute of Developmental and Regenerative Medicine, Department of Physiology, Anatomy and Genetics, University of Oxford, Oxford, UK. Department of Medical Laboratories, College of Applied Medical Sciences, Quassim University, Saudi Arabia (HO Alharbi).

## Abstract

The three striatins (STRN, STRN3, STRN4) form the core of ***<u>STR</u>***iatin-***I***nteracting ***<u>P</u>***hosphatase and ***<u>K</u>***inase (STRIPAK) complexes. These place protein phosphatase 2A (PP2A) in proximity to protein kinases thereby restraining kinase activity and regulating key cellular processes. Our aim was to establish if striatins play a significant role in cardiac remodelling associated with cardiac hypertrophy and heart failure. All striatins were expressed in control human hearts, with upregulation of STRN and STRN3 in failing hearts. We used mice with global heterozygote gene deletion to assess the roles of STRN and STRN3 in cardiac remodelling induced by angiotensin II (AngII; 7 days). Using echocardiography, we detected no differences in baseline cardiac function or dimensions in STRN^+/-^ or STRN3^+/-^ male mice (8 weeks) compared with wild-type littermates. Heterozygous gene deletion did not affect cardiac function in mice treated with AngII, but the increase in left ventricle mass induced by AngII was inhibited in STRN^+/-^ (but not STRN3^+/-^) mice. Histological staining indicated that cardiomyocyte hypertrophy was inhibited. To assess the role of STRN in cardiomyocytes, we converted the STRN knockout line for inducible cardiomyocyte-specific gene deletion. There was no effect of cardiomyocyte STRN knockout on cardiac function or dimensions, but the increase in left ventricle mass induced by AngII was inhibited. This resulted from inhibition of cardiomyocyte hypertrophy and cardiac fibrosis. The data indicate that cardiomyocyte striatin is required for early remodelling of the heart by AngII and identify the striatin-based STRIPAK system as a signalling paradigm in the development of pathological cardiac hypertrophy.

**Clinical perspectives:**

- **Background.** Striatins form the core of ***<u>STR</u>***iatin-***I***nteracting ***<u>P</u>***hosphatase ***<u>A</u>***nd ***<u>K</u>***inase (STRIPAK) complexes that regulate crucial cellular processes such as those associated with heart failure.
- **Summary.** The three striatins are expressed in human hearts, with upregulation of STRN and STRN3 in failing hearts, whilst studies in mice indicate that STRN is required in cardiomyocytes for early remodelling of the hypertensive heart.
- **Potential significance of results to human health and disease.** STRN-based STRIPAKs represent a novel signalling paradigm in the development of pathological cardiac hypertrophy, and modulating this system may provide therapeutic options for managing the cardiac effects of hypertensive heart disease.

## Introduction

The development of heart failure is associated with significant morbidity and a poor prognosis despite optimal medical therapy. It affects millions of people worldwide [1]. A leading cause of heart failure is hypertension that affects ∼30% of all adults [1, 2]. Whilst the identification of patients suffering from hypertension and their management has significantly improved in recent years, a number of patients present acutely with end-organ damage or progress to complications of hypertension in spite of maximal therapy. Broader therapeutic options for individual patients to manage hypertensive heart disease are clearly needed, but this requires greater understanding of the underlying mechanisms. Striatin (STRN) is associated with salt-dependent hypertension in mice. Thus, mice that are heterozygotic for STRN gene deletion have a similar blood pressure profile to wild-type littermates when provided with a low salt diet, but have a greater increase in blood pressure with higher dietary salt [3]. This probably results from effects in endothelial cells and enhanced vasoconstriction [4]. These mice also have increased renal damage in response to aldosterone [5]. Further evidence for a role for STRN in heart failure comes from boxer dogs with arrhythmogenic right ventricular cardiomyopathy (ARVC) and heart failure, resulting from an 8 bp deletion in the 3’UTR of STRN and reduced STRN expression [6, 7]. In humans, SNPs in the STRN gene are linked to blood pressure regulation [3, 8, 9], PR/QRS interval [10, 11], hypertrophic cardiomyopathy [12] and heart failure [13]. SNPs in a second isoform, striatin 3 (STRN3, also known as SG2NA) are also linked to hypertension [14]. STRN, STRN3 and the third isoform, STRN4, are all expressed in the heart. Understanding the roles of these proteins may increase therapeutic options in patients with hypertension to both reduce the risk and to manage the development of heart failure complications.

The heart mainly contains contractile cardiomyocytes, endothelial cells in the capillary network and fibroblasts producing extracellular matrix [15]. Since adult mammalian cardiomyocytes are terminally-differentiated, they respond to increased cardiac workload (e.g. resulting from hypertension) with hypertrophic growth to increase contractile function [16]. The increase in cell size is associated with increased contractile apparatus and changes in gene expression (e.g. upregulation of *Myh7* and *Nppb* mRNAs).

Cardiomyocytes may undergo necrosis or programmed cell death in situations of extreme or prolonged stress, compromising cardiac function and leading to heart failure [17]. Loss of endothelial cells and capillary rarefaction is another feature of developing heart failure [18], along with increased fibrosis in the myocardium (interstitial fibrosis) that compromises cardiac function by increasing ventricular wall stiffness, and around the arterioles (perivascular fibrosis) [19, 20]. Fibrosis is associated with increased numbers of myofibroblasts which may derive from activation of resident fibroblasts or other cardiac cells (e.g. endothelial cells may undergo endothelial to mesenchymal transition and increase numbers of myofibroblasts) [21, 22].

Cellular changes associated with cardiac remodelling are regulated by protein phosphorylation/dephosphorylation. Whilst much is known about protein kinase signalling cascades, specific roles of phosphatases and dephosphorylation are less well understood. The Ser-/Thr-phosphatase PP2A is highly abundant and ubiquitously expressed [23]. It is formed of one of two catalytic subunits (PP2A_C_), one of two regulatory subunits (PP2A_A_) and one of many targeting “B” subunits that direct PP2A to its substrates. Striatins form a class of B subunits (B’’’) for PP2A, but also interact with protein kinases, particularly those of the Germinal Centre Kinase (GCK) family [24]. Because of this, striatin-based complexes have been termed ***<u>STR</u>***iatin-***I***nteracting ***<u>P</u>***hosphatase and ***<u>K</u>***inase (STRIPAK) complexes [25, 26]. This system places GCKs in close proximity to PP2A which maintains the kinase in a dephosphorylated (inactive) state. Inhibition of PP2A or removal of the phosphatase or kinase results in kinase activation, most probably through autophosphorylation. Striatins have an N-terminal domain that binds caveolins, potentially directing them to the plasma membrane and a Ca^2+^/calmodulin-binding domain. A coiled-coil domain and a C-terminal WD repeats facilitate binding of other proteins. Interactome studies have identified many proteins in STRIPAK complexes, some of which probably direct different complexes to different subcellular targets and/or subdomains [27–29]. Striatins, GCKs and STRIPAKs regulate a diverse array of cellular processes including cell survival/proliferation/migration, key features of cardiac remodelling. As described above, STRN is linked to various forms of heart failure and conductance irregularities. Consistent with the latter, STRN is of significant importance in composite junctions between cells, and immunostaining experiments place STRN at intercalated discs between cardiomyocytes, potentially regulating ion fluxes between cells [6, 30].

Our hypothesis is that STRIPAKs play a significant role in cardiac remodelling associated with developing heart failure. Here, we show that *STRN*, *STRN3* and *STRN4* are all dysregulated in human failing hearts compared with normal controls, with upregulation of STRN and STRN3. Our data in mouse models with heterozygote knockout of STRN and STRN3 identified STRN, but not STRN3, as a potential mediator of the early phase of cardiac remodelling (i.e. prior to heart failure development and cardiac dysfunction) induced by developing hypertension in mice resulting from angiotensin II (AngII) infusion. Further studies using mice with inducible cardiomyocyte-specific STRN deletion confirmed that cardiomyocyte striatin plays a key role in this early remodelling phase.

## Methods

### Ethics statement

#### Human heart samples

Human heart samples were from the University of Pittsburgh, U.S.A. Failing human heart samples were from patients who consented to a protocol reviewed and approved by the University of Pittsburgh Institutional Review Board. Non-failing heart samples were collected under University of Pittsburgh CORID #451 (Committee for Oversight of Research and Clinical Training Involving Decedents) and with consent being obtained by the local Organ Procurement Organization (OPO), CORE (Center for Organ Recovery and Education).

#### Mouse studies

Mice were housed at the BioResource Unit at University of Reading (colonies for STRN and STRN3 global knockout) or St. George’s University of London (colonies for cardiomyocyte-specific deletion of STRN), both UK registered with a Home Office certificate of designation. Procedures were performed in accordance with UK regulations and the European Parliament Directive 2010/63/EU for animal experiments. All work was undertaken in accordance with local institutional animal care committee procedures at the University of Reading and the U.K. Animals (Scientific Procedures) Act 1986. Studies were conducted under Project Licences 70/8248, 70/8249 and P8BAB0744.

### Studies of striatin isoforms in human hearts

mRNA expression of *STRN* (ENSG00000115808), *STRN3* (ENSG00000196792) and *STRN4* (ENSG00000090372) was determined using a previously published RNASeq dataset derived from left ventricular samples of patients with end-stage dilated cardiomyopathy (n=97) taken at the time of transplantation or left ventricular assist device implantation, compared with non-diseased controls (n=108) [31]. Differential expression analysis was performed with DESeq2 (V*1.18.1*, Wald test) [32].

Human heart samples used in this study were previously used to study RAF kinases [33]. Transmural tissue at the level of the anterior papillary muscle was collected at the time of cardiac transplantation from the left ventricle of end-stage heart failure patients. Samples were collected in the operating room and transported in ice-cold St. Thomas’ cardioplegia solution, flash frozen within 20 minutes of excision, and stored at -80°C prior to utilization.

Control left ventricular tissues were collected from hearts that were rejected for transplant for varying reasons. Tissues were collected and stored in a similar manner as the failing hearts, with between 20-45 minutes of time elapsing between cross-clamp and freezing of the tissue. Hearts were ground to powder under liquid N_2_, and samples taken for RNA and protein preparation as described below.

### Animal husbandry and randomisation

Housing conditions were as described in [33, 34]. Animals were checked daily and breeding was conducted with mice between 6 weeks and 8 months with a maximum of 6 litters per female. Mice undergoing procedures were monitored using a score sheet and routinely culled if they reached a predefined endpoint agreed with the Named Veterinary Surgeon. Weights were taken before, during and at the end of the procedures. Mouse weights from the start and end of procedures are provided in **Supplementary Table S1**. These studies used only male mice because of the intention to convert the line for conditional gene deletion using tamoxifen (see below), an approach which has not been fully characterised for female mice. Furthermore, our recent studies of inducible deletion of BRAF indicated that males and females responded very differently to this regime [35]. Mice were allocated to specific groups on a random basis with randomisation performed independently of the individual leading the experiment. Four mice receiving AngII that died and all data for these were excluded from analysis (one wild-type from the STRN colony that died on day 2, two STRN^fl/fl^/Cre^MCM/-^ mice treated with corn-oil that died on day 7 and one STRN^fl/fl^/Cre^MCM/-^ mouse treated with tamoxifen that died on day 7). Post-mortem analysis showed rupture of a major blood vessel in all cases. Otherwise, no mice were excluded after randomisation. Individuals conducting the studies were not blinded to experimental conditions for welfare monitoring purposes. Data and sample analysis (e.g. echocardiography, histology) was performed by individuals who were blinded to intervention.

### Mouse lines and gene deletion strategy

Mice for STRN or STRN3 knockout were from the Knockout Mouse Project (KOMP). Both lines (“Knockout first” STRN^tm1a(KOMP)WTsi^ and STRN3^tm1a(KOMP)WTsi^) were on a C57Bl/6N background and, following resuscitation, were backcrossed with C57Bl/6J mice (Charles River Laboratories) for at least eight generations prior to experimentation and sperm preservation (at the Mary Lyon Centre, MRC Harwell, UK). Colonies were maintained as heterozygotes with ongoing breeding with C57Bl/6J mice to generate heterozygote and wild-type (WT) littermates for experiments. (N.B. Global homozygous knockout of any striatin isoform is embryonic lethal).

*Myh6*-MERCreMER mice expressing tamoxifen-inducible Cre recombinase under control of a mouse *Myh6* promoter [Tg(Myh6-cre)1Jmk/J, strain no. 009074] [36] were from Jackson Laboratories, imported into the UK and transported to St. George’s University of London for breeding in-house. These mice are on a C57Bl/6J background. Mice for cardiomyocyte-specific knockout of STRN were derived from the sperm banked at the Mary Lyon Centre from the STRN mice. The line was resuscitated and allele conversion using FLP recombinase to generate the conditional ready floxed line was performed by the Mary Lyon Centre (MRC Harwell). Heterozygous floxed STRN (STRN^WT/fl^) mice were transported to St. George’s University of London and backcrossed onto a C57Bl/6J background for 4 generations maintaining the line as heterozygotes before generating the homozygote line (STRN^fl/fl^). They were then crossed with homozygous Cre (Cre^+/+^) mice to generate mice that were heterozygous STRN and hemizygous for Cre (STRN^WT/fl^/Cre^+/-^); these were used to generate double homozygotes (STRN^fl/fl^/Cre^+/+^). STRN^fl/fl^ mice were bred with STRN^fl/fl^/Cre^+/+^ mice to generate mice hemizyogous for Cre and homozygous for floxed STRN (STRN^fl/fl^/Cre^+/^).

Tamoxifen was dissolved in 0.25 ml ethanol which was then mixed with 4.75 ml corn oil. Male mice (8-9 wks) were treated with a single dose of tamoxifen (40 mg/kg i.p.; Sigma-Aldrich) to induce recombination or corn-oil vehicle as a control at -4 days relative to mini-pump implantation (see below). Our previous studies of mice hemizygous for Cre^MCM/-^ treated in this way demonstrated that this is sufficient to induce recombination but has no overt effect on cardiac function or dimensions at baseline or on AngII-induced cardiac hypertrophy [33, 35].

### Genotyping and confirmation of recombination

Ear notches were taken for identification using a 0.5 mm ear punch and used for genotyping. For confirmation of recombination in the heart, hearts and kidneys were collected from mice treated with tamoxifen or corn-oil vehicle, the tissues were ground to powder under liquid N_2_ and samples were taken. DNA was extracted using Purelink genomic DNA (gDNA) mini-kits (Invitrogen) according to the manufacturer’s instructions. gDNA was purified through Purelink spin columns and eluted in 30 µl of elution buffer. PCR amplification used GoTaq Hot Start Polymerase (Promega). PCR conditions were 95°C for 3 min, followed by up to 35 cycles of 95°C denaturations for 30 s, 30 s annealing, elongation at 72°C for 30 s, followed by a 7-minute 72°C final extension. Details of primers and conditions are in **Supplementary Table S3**. PCR products were separated using gel electrophoresis (25 minutes, 80 V) on 2% (w/v) agarose gels and visualised under UV light.

### AngII-induced cardiac hypertrophy

Alzet osmotic minipumps (supplied by Charles River Laboratories) were used for continuous delivery of 0.8 mg/kg/d AngII or vehicle for 7 d. Mice were anaesthetised in an induction chamber using vaporised 5% isoflurane in a constant oxygen supply of 2 l/min. Anaesthesia was maintained at 2.5% isoflurane using a nose cone. Mice were positioned on a heated mat in the prone position. Buprenorphine (Vetergesic, Ceva Animal Health Ltd.) (0.05 mg/kg, diluted in sterile PBS) was administered subcutaneously for analgesia. The fur covering the mid-scapular region was removed using an electric razor and the area was sterilised with HIBISCRUB^®^ (VioVet). Under aseptic conditions, a 2 cm incision was made at the mid-scapular region and blunt dissection generated a pocket towards the lower-left flank of the mouse for the minipump to be inserted. The wound was closed with two simple interrupted sutures using polypropylene 4-0 thread (Prolene, Ethicon) and then sterilised with HIBISCRUB^®^. Mice were recovered singly and returned to a clean cage once fully recovered.

### Mouse echocardiography

Echocardiography was performed using a high-frequency ultrasound system (Vevo 2100^TM^, Visualsonics) equipped with a 38 MHz MS400 transducer. Baseline echocardiograms were collected at 8 weeks (prior to tamoxifen treatment and/or minipump implantation) with additional scans taken at the end of the study. Mice were anaesthetised in an induction chamber using vaporised 5% isoflurane in a constant oxygen supply of 1 l/min. Anaesthesia was maintained with 1.5% isoflurane using a nose cone. Mice were positioned on a heating physiological monitoring stage in a supine position. Heart rate, respiration rate and body temperature were monitored. Chest fur was removed with an electric razor and hair removal cream. Pre-warmed ultrasound gel was applied to the chest as a coupling medium for the transducer. The transducer was orientated and lowered into the ultrasound gel until a clear image was centralised on the monitor. Imaging was completed within a maximum time of 30 min, and usually within 15 min. Mice were recovered singly and transferred to the home cage once fully recovered. Cardiac function and global longitudinal strain were measured from B-mode long axis images using VevoStrain software for speckle tracking. B-mode images of the ascending aorta were also captured. At the end of the experiment, whilst still under anaesthesia, mice were culled by cervical dislocation with severance of the femoral artery to ensure cessation of life. Hearts were excised quickly, washed in PBS, dried and snap-frozen in liquid N_2_ or fixed for histology.

### Histology and analysis

Histological sections for the global STRN and STRN3 knockout mouse studies were prepared and stained by HistologiX Limited. Sections for the cardiomyocyte-specific STRN knockout study were prepared and stained at St. George’s University of London (as described in [37]). Haemotoxylin and eosin staining was used for analysis of myocyte cross-sectional area. Cells around the periphery of the left ventricle (excluding epicardial layer) were chosen at random (ensuring that the cells were in cross section and with a clear, rounded nucleus) and outline traced using NDP.view2 software (Hamamatsu). Up to 30 cells were measured per section by a single independent assessor and the mean value taken for each mouse. To assess interstitial fibrosis, sections were stained with Masson’s trichrome or picrosirius red and analysis used Image-J as in [37]. The collagen fraction was calculated as the ratio between the sum of the total area of fibrosis (blue colour for Masson’s trichrome, red colour for picrosirius red) to the sum of the total tissue area (including the myocyte area) for the entire image and expressed as a percentage. For perivascular fibrosis (because there was not a constant number of vessels apparent in each section), picrosirius red staining was used and the whole section was scored for perivascular fibrosis around arterioles (identified by a clear elastic layer). Values were from 1 (negligible increase in fibrosis around any vessel), through to 4 (extensive fibrosis around multiple vessels, penetrating into the myocardium).

### RNA preparation and qPCR

Heart powders (10-15 mg) were weighed into safelock Eppendorf tubes and kept on dry ice. RNA Bee (AMS Biotechnology Ltd) was added (1 ml) and the samples homogenised on ice using a pestle. RNA was prepared according to the manufacturer’s instructions and dissolved in nuclease-free water. The purity was assessed from the A_260_/A_280_ measured using an Implen NanoPhotometer (values were 1.8–2.0) and concentrations determined from the A_260_. Quantitative PCR (qPCR) analysis was performed as described in [38]. Total RNA was reverse transcribed to cDNA using High Capacity cDNA Reverse Transcription Kits with random primers (Applied Biosystems). qPCR was performed using a StepOnePlus Real-Time PCR system (ThermoFisher Scientific) using 1/40 of the cDNA produced. Optical 96-well reaction plates were used with iTaq Universal SYBR Green Supermix (Bio-Rad Laboratories Inc.) according to the manufacturer’s instructions. See **Supplementary Table S4** for primer sequences. Results were normalized to *GAPDH*, and relative quantification was obtained using the ΔCt (threshold cycle) method; relative expression was calculated as 2^−ΔΔCt^, and normalised as indicated in the Figure Legends.

### Immunoblotting

Heart powders (15-20 mg) were homogenised in 6 vol extraction buffer [20 mM Tris pH 7.5, 1 mM EDTA, 10% (v/v) glycerol, 1% (v/v) Triton X-100, 100 mM KCl, 5 mM NaF, 0.2 mM Na_3_VO_4_, 5 mM MgCl_2_, 0.05% (v/v) 2-mercaptoethanol, 10 mM benzamidine, 0.2 mM leupeptin, 0.01 mM trans-epoxy succinyl-L-leucylamido-(4-guanidino)butane, 0.3 mM phenylmethylsulphonyl fluoride, 4 µM microcystin]. Samples were extracted on ice with intermittent vortex mixing (10 min), then centrifuged (10,000 × g, 10 min, 4°C) to pellet insoluble material. The supernatants were removed, a sample was taken for protein assay and the rest boiled with 0.33 vol sample buffer (300 mM Tris-HCl pH 6.8, 10% (w/v) SDS, 13% (v/v) glycerol, 130 mM dithiothreitol, 0.2% (w/v) bromophenol blue). Protein concentrations were determined by BioRad Bradford assay using a 1/5 dilution (v/v) in H_2_O of the dye reagent concentrate and BSA standards.

Proteins (100 µg for human heart samples, 40 µg for rat and mouse heart samples), were separated by SDS-PAGE (200 V) using 8% (for striatin isoforms), or 12% (GAPDH) polyacrylamide resolving gels with 6% stacking gels until the dye front reached the bottom of the gel (∼50 min). Proteins were transferred electrophoretically to nitrocellulose using a BioRad semi-dry transfer cell (10 V, 60 min). Non-specific binding sites were blocked (15 min) with 5% (w/v) non-fat milk powder in Tris-buffered saline (20 mM Tris-HCl pH 7.5, 137 mM NaCl) containing 0.1% (v/v) Tween 20 (TBST). Blots were incubated with primary antibodies in TBST containing 5% (w/v) BSA (overnight, 4°C), then washed with TBST (3 × 5 min, 21°C), incubated with horseradish peroxidase-conjugated secondary antibodies in TBST containing 1% (w/v) non-fat milk powder (60 min, 21°C) and then washed again in TBST (3 × 5 min, 21°C). Rabbit polyclonal antibodies to STRN and STRN4 were from Novus Biologicals Ltd (STRN: catalogue number NB110-74571; STRN4: catalogue no. NBP2-36537) and were used at 1/1000 dilution. Goat polyclonal antibodies to STRN3 (SG2NA) were from Santa Cruz Biotechnology Inc (catalogue no. E1704). Rabbit polyclonal antibodies to GAPDH were from Cell Signaling Technologies (catalogue no. 14C10). All primary antibodies were used at 1/1000 dilutions. Horseradish-peroxidase-conjugated goat anti-rabbit immunoglobulins (catalogue no. P0448) and rabbit anti-goat immunoglobulins (catalogue no. P0449) were from Dako (supplied by Agilent) and were used at 1/5000 dilution. Bands were detected by enhanced chemiluminescence using ECL Prime with visualisation using an ImageQuant LAS4000 system (Cytiva). ImageQuant TL 8.1 software (GE Healthcare) was used for densitometric analysis. Raw values for phosphorylated kinases were normalised to the total kinase. Values for all samples were normalised to the mean of the controls.

### Image processing and statistical analysis

Images were exported from the original software as .tif or .jpg files and cropped for presentation with Adobe Photoshop CC maintaining the original relative proportions. Data analysis used Microsoft Excel and GraphPad Prism 9. Statistical analysis was performed using GraphPad Prism 9. A Grubb’s outlier test was applied to the data, and outliers excluded from the analysis. Statistical significance was determined using two-tailed unpaired Mann-Whitney tests, or two-tailed one-way or two-way ANOVA as indicated in the Figure Legends. A Holm-Sidak’s multiple comparison test was used in combination with ANOVA. Graphs were plotted with GraphPad Prism 9. Specific p values are provided with significance levels of p<0.05 in bold type.

## Results

### Striatin and striatin 3 are upregulated in human failing hearts

To assess which of the striatin isoforms is most likely to promote human heart failure, we mined an RNASeq database of heart samples from patients with dilated cardiomyopathy (n=97) compared with normal controls (n=108) [31]. *STRN*, *STRN3* and *STRN4* transcripts were readily detected, with a rank order of expression of *STRN4*>*STRN*>*STRN3* in control samples (**Figure 1A**). Expression of STRN4 declined in DCM hearts whilst expression of *STRN* and *STRN3* increased. We also assessed expression of striatins in samples from 12 patients with heart failure of mixed non-ischaemic aetiology compared with normal controls (previously reported in [33]). These showed a significant increase in only *STRN3* mRNA expression in heart failure samples (**Figure 1B**), although protein expression of all striatins was significantly increased in failing hearts (**Figure 1C-D**).

**Figure 1.**
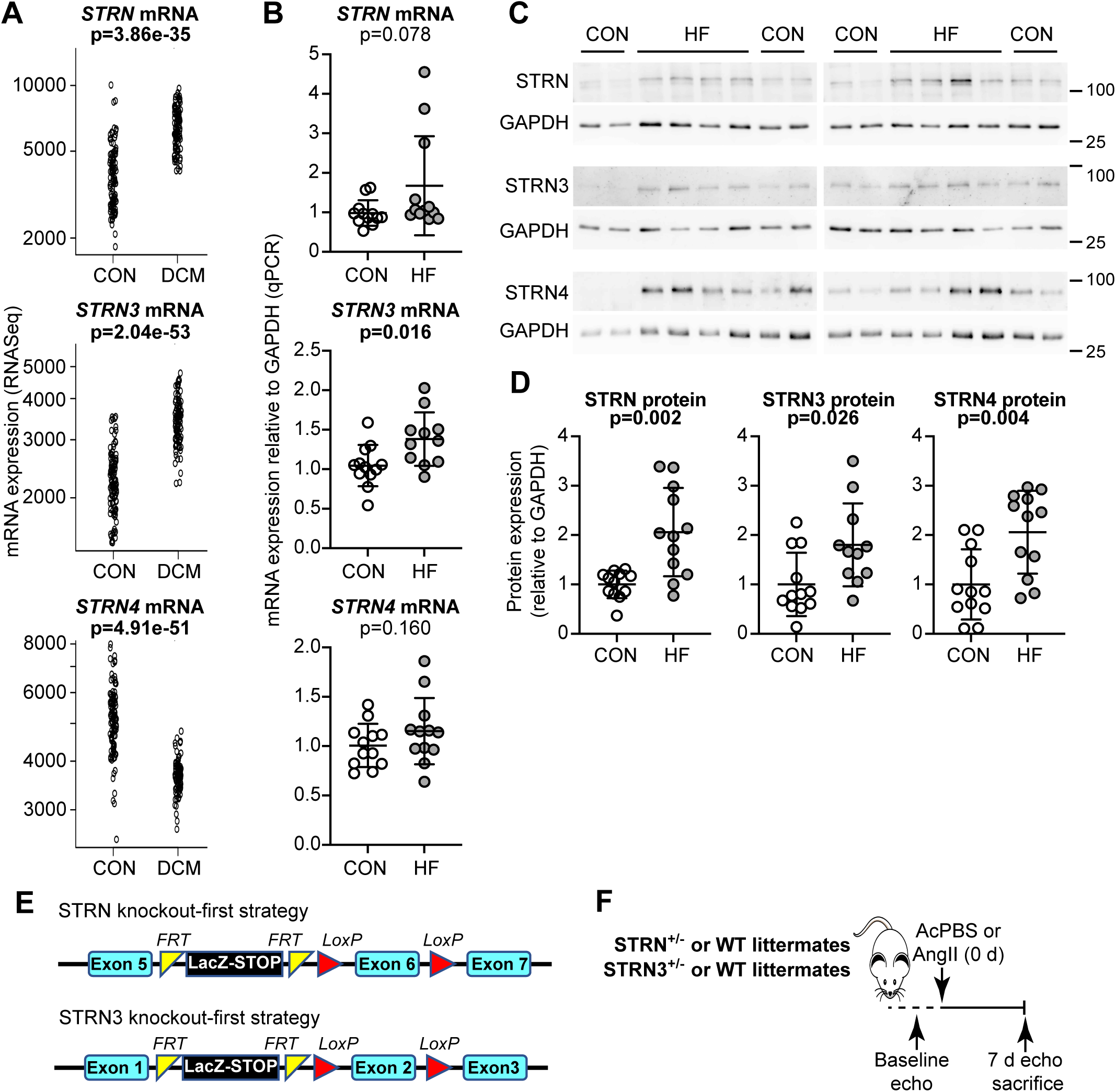
Expression of striatin isoforms in human failing hearts and strategy for STRN or STRN3 knockout in mice. **A,** Data for mRNA expression of striatin isoforms in human hearts were from an RNASeq database of patients with dilated cardiomyopathy (DCM, n=97) and normal controls (CON, n=108). Data for individual samples are shown with adjusted p values. **B-D**, Samples from control hearts (CON) or explanted hearts from patients with heart failure (HF) were used to prepare RNA for qPCR analysis (**B**) or protein for immunoblots (**C-D**). Representative immunoblots (100 µg protein per lane) are in **C** with densitometric analysis in **D**. Individual values are shown with means ± SD. Results are relative to GAPDH and normalised to the mean of the CON hearts. Mann-Whitney tests were used for statistical analysis. **E,** “Knockout-first” strategy for global deletion of STRN or STRN3 in mice involved positioning of a STOP cassette flanked by FRT sites upstream of a critical exon that was also flanked with LoxP sites. **F,** Experimental approach for assessment of effects of STRN or STRN3 deletion on cardiac function. Homozygous global knockout of STRN or STRN3 is embryonic lethal, so heterozygote STRN^+/-^ or STRN3^+/-^ mice were used in comparison with wild-type (WT) littermates from each colony. Following baseline echocardiography (echo), minipumps were implanted for delivery of acidified PBS vehicle (AcPBS) or 0.8 mg/kg/d angiotensin II (AngII). Following echocardiography at 7 d, mice were sacrificed.

### Heterozygous knockout of STRN in mice, but not STRN3, reduces cardiac hypertrophy induced by AngII

Since STRN and STRN3 were upregulated in DCM and human failing hearts of other aetiology, we hypothesized that these isoforms were most likely to be involved in developing cardiac pathology. To assess this, we used commercially-available knockout-first mice, engineered with a removable STOP cassette to permit conversion for conditional gene deletion (**Figure 1E**). Homozygous global deletion of any striatin isoform is embryonic lethal, so we assessed the effects of heterozygous gene deletion (STRN^+/-^ and STRN3^+/-^), comparing responses with wild-type littermates. We studied male mice taking baseline echocardiograms at 8 weeks, and detected no differences in cardiac function or dimensions in STRN^+/-^ or STRN3^+/-^ mice compared with the wild-types (**Supplementary Table S4**). Mice were then treated for 7 days with acidified PBS vehicle or with 0.8 mg/kg/d AngII using osmotic minipumps (**Figure 1F**). This “slow-pressor” dose gradually induces hypertension over 7-14 days [39–41].

STRN protein (upper band on immunoblots at ∼110 kDa) was reduced in hearts from STRN^+/-^ mice (**Figure 2A-B**). This did not affect expression of STRN4, but there was a small, non-significant (p=0.059) increase in STRN3 suggesting there could be some compensation in some animals. AngII increased expression of STRN in wild-type littermates from the STRN colony, and this was accompanied by a significant increase in STRN3.

**Figure 2.**
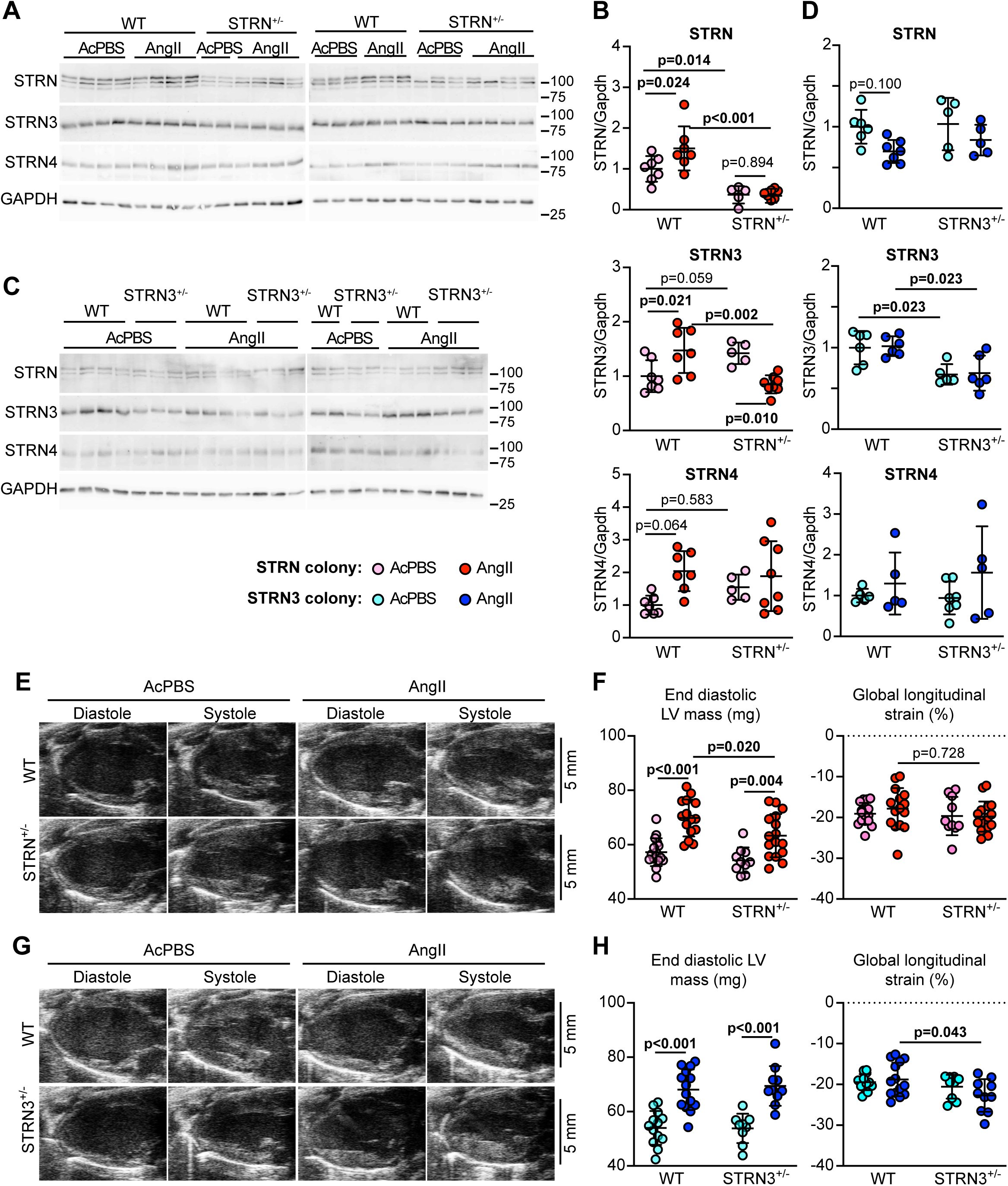
Heterozygous global deletion of STRN, but not STRN3, compromises the hypertrophic response to AngII. Male mice (8 wks) heterozygote for STRN or STRN3 knockout (STRN^+/-^ or STRN3^+/-^) and wild-type (WT) littermates from each colony were treated with acidified PBS (AcPBS) vehicle or AngII (0.8 mg/kg/d). Data for STRN colony mice are in **A, B, E** and **F**; data for STRN3 colony mice are in **C, D, G** and **H**. **A-D,** Heart powders were used for immunoblotting (40 µg protein per lane). Representative immunoblots of the striatin isoforms and GAPDH (**A, C**) are shown with **d**ensitometric analysis (**B, D**). Results are relative to GAPDH and normalised to the means for WT mice treated with AcPBS. N.B. The upper band of the STRN blot used for densitometry correlates with the predicted molecular weight of STRN protein (110k Da). **E-H**, Echocardiography of mouse hearts taken at 7 d with representative long-axis images (**E, G**) and estimated end diastolic left ventricle (LV) mass and global longitudinal strain (**F, H**). Individual datapoints are plotted with means ± SD. Statistical analysis used 2-way ANOVA with Holm-Sidak’s post-test.

STRN3 was reduced in hearts from STRN3^+/-^ mice with no significant change in expression of STRN or STRN4 (**Figure 2C-D**). In contrast to the STRN colony, we did not detect an increase in any striatin isoform with AngII in hearts from wild-type mice from the STRN3 colony, presumably reflecting differences in the background strain despite extensive backcrossing onto the same C57Bl/6J background.

We assessed the effects of AngII on the heart using echocardiography (**Figure 2E-H; Supplementary Table S5**). There were no significant changes in cardiac function parameters (heart rate, ejection fraction, fractional shortening, cardiac output) or in estimated left ventricule (LV) volume. However, AngII induced a significant increase in estimated LV mass in wild-type mice from each of the STRN and STRN3 colonies and this was significantly reduced in STRN^+/-^ mice compared with their wild-type littermates (**Figure 2F**). This was not associated with any differences in global longitudinal strain between wild-type and STRN^+/-^ mice. Interestingly, STRN3^+/-^ mice had a similar increase in LV mass as their wild-type littermates, but there was an increase in global longitudinal strain (**Figure 2H**), suggesting there was an effect on longitudinal contraction of the heart.

Heart sections were stained with haemotoxylin and eosin and the cross-sectional area of myocytes at the periphery of the LV measured. AngII increased cardiomyocyte cross-sectional area (indicative of cardiomyocyte hypertrophy) in wild-type mice from both STRN and STRN3 colonies (**Figure 3A-D**), and increased hypertrophic gene marker expression (**Figure 3E-F**). The AngII-induced increase in cross-sectional area and expression of *Myh7* mRNA (though not *Nppb*) was reduced in STRN^+/-^ (but not STRN3^+/-^) mice compared with their wild-type littermates. Heart sections were stained with picrosirius red to assess cardiac fibrosis. At this 7 day time point, AngII induced only a small, non-significant increase in interstitial fibrosis and this was most commonly detected in the area at the junction between the interventricular septum and the ventricular wall. However, there was a greater, significant increase in perivascular fibrosis. The degree of fibrosis induced by AngII was similar in STRN^+/-^, STRN3^+/-^ and their wild-type littermates (**Figure 4A-D**). AngII also upregulated mRNAs encoding fibrotic genes but, again, this was not significantly different in STRN^+/-^ or STRN3^+/-^ mouse hearts compared with their wild-type littermates (**Figure 4E-F**).

**Figure 3.**
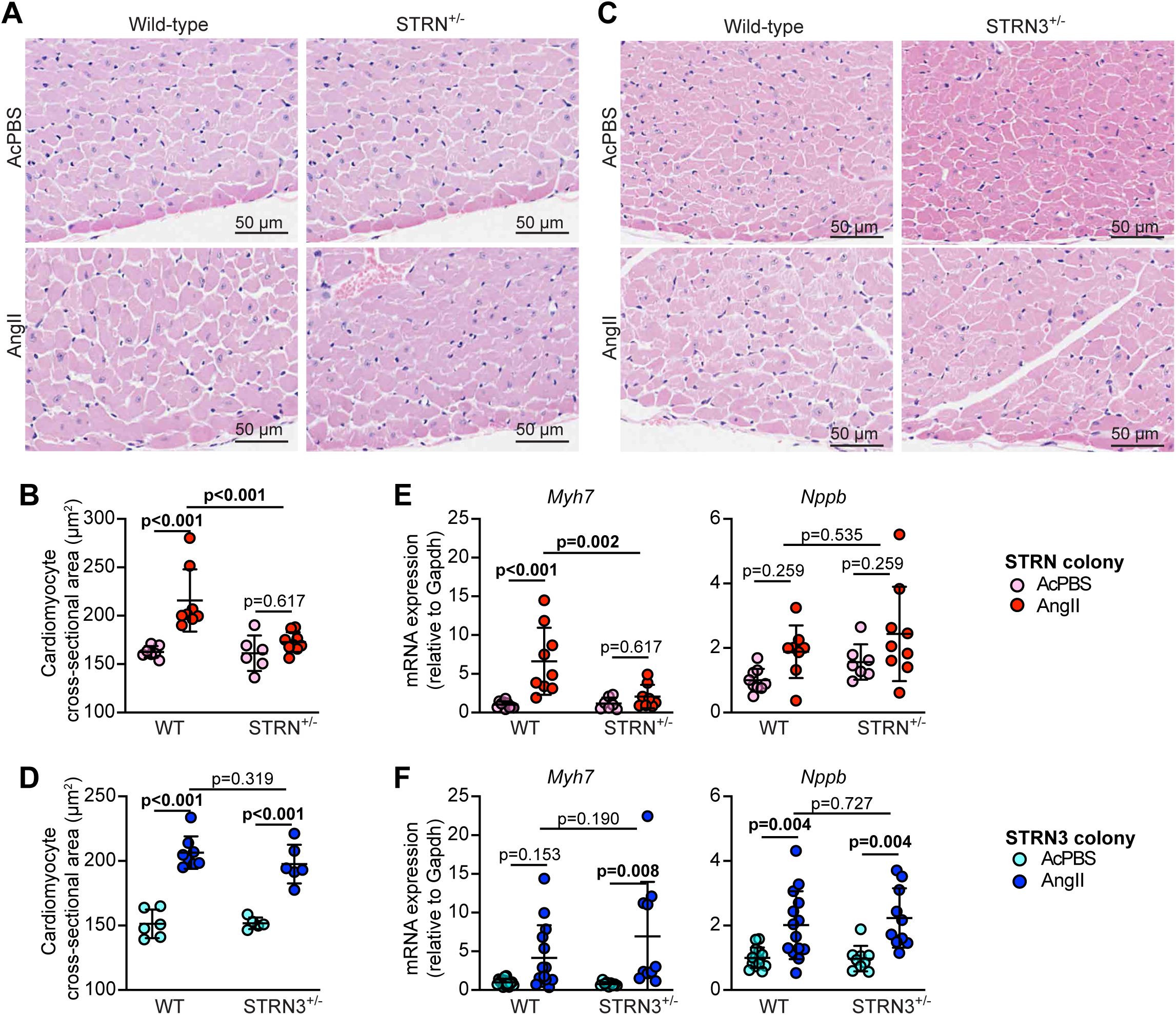
Heterozygous global deletion of STRN, but not STRN3 reduces cardiomyocyte hypertrophy and the increase in expression of *Myh7* induced by AngII. 8 wk male STRN^+/-^ (**A, B** and **E**) and STRN3^+/-^ (**C, D** and **F**) mice, plus their respective wild-type (WT) littermates were treated with acidified PBS (AcPBS) vehicle or 0.8 mg/kg/d AngII (7 d). Hearts were fixed and sections stained with haemotoxylin and eosin. Representative images (**A, C**) show areas from the outer perimeter of the left ventricular wall opposite the interventricular septum. **B, D,** Cardiomyocyte cross-sectional areas are shown. **E-F,** RNA was extracted from mouse heart powders and analysed by qPCR for *Myh7* and *Nppb*. Individual datapoints are plotted with means ± SD. Results are relative to GAPDH and normalised to the means for WT mice treated with AcPBS. Statistical analysis used 2-way ANOVA with Holm-Sidak’s post-test.

**Figure 4.**
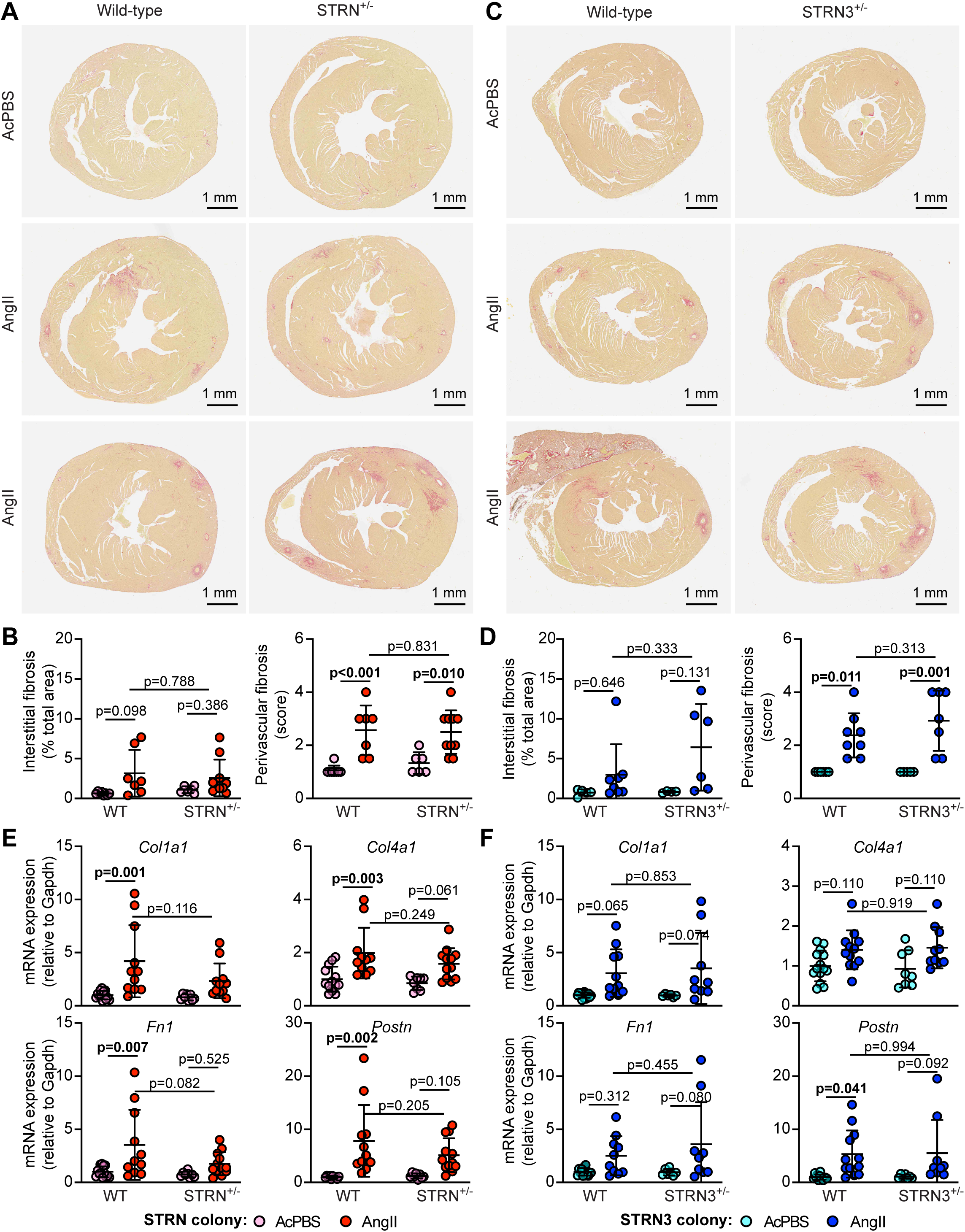
Heterozygous knockout of STRN, or STRN3 does not reduce cardiac fibrosis induced by AngII. 8 wk male STRN^+/-^ (**A, B** and **E**) and STRN3^+/-^ (**C, D** and **F**) mice, plus their respective wild-type (WT) littermates were treated with acidified PBS (AcPBS) vehicle or 0.8 mg/kg/d AngII (7 d). Hearts were fixed and sections stained with picrosirius red. Representative images (**A**, **C**) are from the short axis of the heart, with middle and lower panels illustrating the range of response for each condition. **B, D,** Interstitial fibrosis was measured using Image J and is presented as the % of the total area (excluding regions around the blood vessels). Perivascular fibrosis was scored (1: negligible increase in fibrosis around any vessel; 2: mild to moderate fibrosis around 1 or more vessels; 3: Significant fibrosis permeating tissue around 1 or more vessels; 4: extensive fibrosis around multiple vessels, penetrating into the myocardium). **E-F,** RNA was extracted from mouse heart powders and analysed by qPCR for fibrosis genes (*Col1a1, Col4a1, Fn1* and *Postn1*). Results are relative to GAPDH and normalised to the means for WT mice treated with AcPBS. Individual datapoints are plotted with means ± SD. Statistical analysis used 2-way ANOVA with Holm-Sidak’s post-test.

Overall, the data indicate that reduction of STRN3 does not substantially affect cardiac hypertrophy induced by AngII, at least over the short term. In contrast, reduction of STRN compromises the cardiac response to AngII, having a clear effect on cardiomyocyte hypertrophy, though not cardiac fibrosis.

### Cardiomyocyte-specific deletion of STRN inhibits AngII-induced cardiac hypertrophy

To determine if the reduced hypertrophic response to AngII in STRN^+/-^ mice was due to reduced expression of STRN in cardiomyocytes rather than other cardiac cells, we converted the STRN knockout-first line for conditional gene deletion using FLP recombinase (**Figure 5A**). These mice were used to generate homozygous floxed STRN mice with a single allele for tamoxifen-inducible Cre under the control of a Myh6 promoter [36]. Male STRN^fl/fl^/Cre^+/-^ mice (8 weeks) were treated with a single dose of tamoxifen (40 mg/kg) to induce recombination, an approach which is not associated with significant cardiotoxicity from the Cre enzyme [33, 35]. Recombination was detected in hearts but not kidneys from STRN^fl/fl^/Cre^+/-^ mice treated with tamoxifen (**Figure 5B**), confirming cardiac-specific gene deletion. Osmotic minipumps were implanted 4 days after tamoxifen treatment (by which time the tamoxifen has been cleared from the body [42]) to deliver acidified PBS vehicle or 0.8 mg/kg/d AngII for 7 days. Immunoblotting confirmed that tamoxifen induced a significant and substantial decrease in STRN expression in the hearts of STRN^fl/fl^/Cre^+/-^ mice, and this was associated with a significant increase in expression of STRN3 (**Figure 5D-E**). STRN4 expression was increased in the hearts of some mice, but the overall difference was not statistically significant. As with global heterozygous STRN^+/-^ mice (**Figure 2A-B**), AngII increased expression of STRN, but there was no increase in the hearts of cardiomyocyte STRN knockout mice (**Figure 5D-E**).

**Figure 5.**
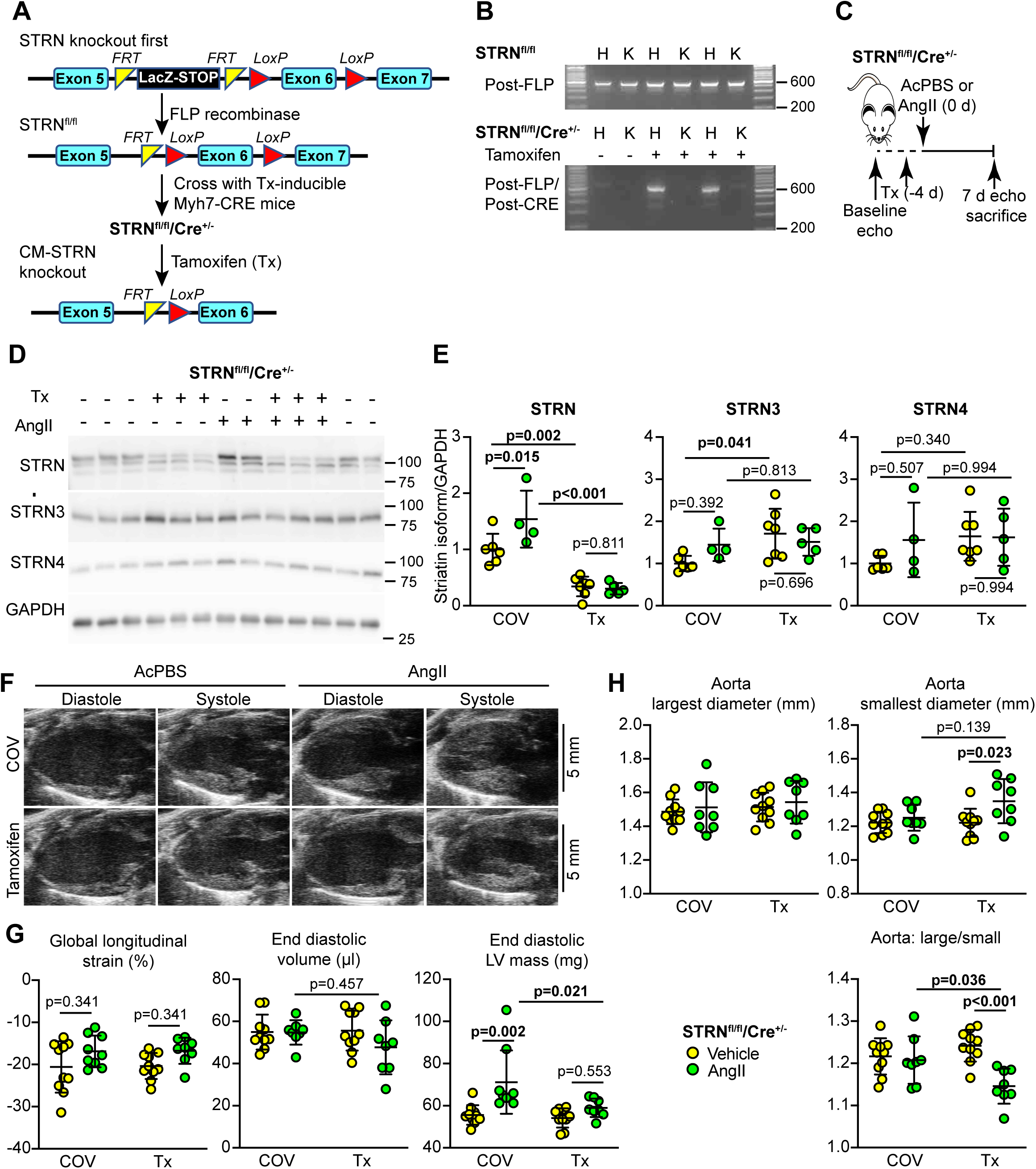
Cardiomyocyte-specific knockout of STRN inhibits cardiac hypertrophy induced by AngII. **A**, Strategy for cardiomyocyte (CM) specific knockout of STRN in mice. STRN knockout first mice were converted to “conditional-ready” using FLP recombinase, removing the STOP cassette between exons 5 and 6, whilst leaving the LoxP sites surrounding exon 6 in place (STRN^fl/fl^ mice). These were bred with mice expressing tamoxifen-(Tx-) activated Cre to generate mice homozygous for floxed striatin and hemizygous for Cre (STRN^fl/fl^/Cre^+/-^) for experiments. Treatment with tamoxifen induced recombination and deletion of exon 6. **B,** Hearts (H) and kidneys (K) from male mice were genotyped to confirm that the mice were conditional-ready (upper panel) and that tamoxifen treatment (40 mg/kg) induced recombination in the heart but not kidney. **C**, Strategy for experiments. Male STRN^fl/fl^/Cre^+/-^ mice (8 wks) were used. Following baseline echocardiography (echo), mice were treated with corn-oil vehicle (COV) or Tx in COV (40 mg/kg; day -4) and minipumps were implanted (day 0) to deliver acidified PBS (AcPBS) vehicle or 0.8 mg/kg/d AngII for 7 days, after which a final echocardiogram was taken before the mice were sacrificed. **D-E**, Heart powders were used for immunoblotting (40 µg protein per lane). Representative immunoblots of the striatin isoforms and GAPDH are in **D**, with densitometric analysis in **E**. Results are relative to GAPDH and normalised to the means for mice treated with vehicle only. The upper band of the STRN blot used for densitometry correlates with the predicted molecular weight of STRN protein. **F-G**, Echocardiography of mouse hearts taken at 7 days with representative long-axis images (**F**) and cardiac dimensions (LV, left ventricle) and longitudinal strain (**G**). **H**, Aortic width measured after the aortic valve at its largest (i.e. immediately following cardiac contraction) and smallest diameter (with cardiac relaxation), with the ratio between the two values. Individual datapoints are plotted with means ± SD. Statistical analysis used 2-way ANOVA with Holm-Sidak’s post-test.

Using echocardiography, we detected no significant differences in cardiac function between any of the conditions (**Supplementary Table S6**). As in the studies of global heterozygous knockout of STRN, AngII did not significantly affect predicted LV volumes, but increased estimated LV mass (**Figure 5F-G**). This was significantly reduced in mice with cardiomyocyte deletion of STRN. To determine if there were likely to be secondary consequences, we measured the width of the aorta after systolic contraction when the aortic diameter is at its largest, compared with the diameter following cardiac relaxation. AngII had no significant effect on either measurement in mice treated with corn-oil vehicle alone. However, with cardiomyocyte deletion of STRN, the width of the aorta following relaxation (i.e. the smallest diameter) was significantly greater in mice treated with AngII than the aortas from the vehicle treated mice (**Figure 5H**). This resulted in a significant decrease in the width ratio, suggesting that the loss of striatin in the heart had a secondary effect on the aorta.

AngII increased cardiomyocyte cross-sectional area and this was significantly reduced with cardiomyocyte knockout of STRN (**Figure 6A-B**). However, hypertrophic gene marker expression (*Myh7* and *Nppb*) was similar with or without tamoxifen treatment (**Figure 6C**) suggesting the cells continued to undergo pathological stress. Unexpectedly and in contrast to the effect of global heterozygous STRN knockout (**Figure 3A-B**), the increase in interstitial and perivascular fibrosis induced by AngII was significantly inhibited with cardiomyocyte STRN knockout and this was associated with reduced expression of fibrotic gene markers (**Figure 6D-F**). We conclude that cardiomyocyte STRN plays an important role in early adaptive remodelling of the heart induced by AngII, with effects at the level of the cardiomyocytes themselves to promote hypertrophic growth and to increase cardiac fibrosis.

**Figure 6.**
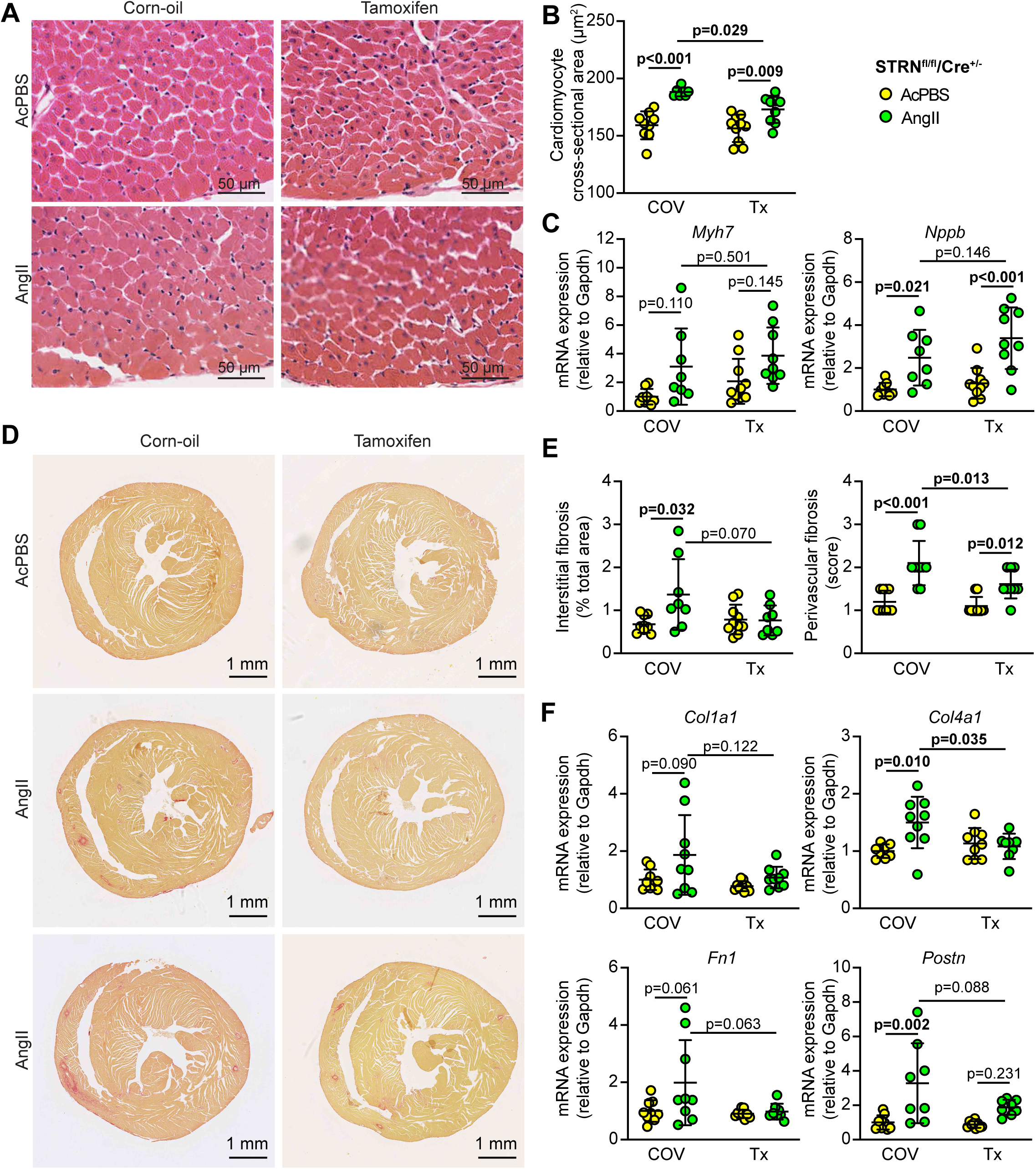
Cardiomyocyte-specific knockout of STRN inhibits the increase in cardiomyocyte cross-sectional area and fibrosis induced induced by AngII. Male STRN^fl/fl^/Cre^+/-^ mice (8 wks) were treated with corn-oil vehicle (COV) or tamoxifen (Tx) (40 mg/kg) and minipumps implanted to deliver acidified PBS (AcPBS) or 0.8 mg/kg/d AngII for 7 days. Hearts were fixed or snap-frozen in liquid N_2_ before grinding to powder. Representative images of sections stained with haemotoxylin and eosin (**A**) show areas from the outer perimeter of the left ventricular wall opposite the interventricular septum. **B,** Cardiomyocyte cross-sectional areas. **C,** RNA extracted from mouse heart powders was analysed by qPCR for *Myh7* and *Nppb*. Results are relative to GAPDH and normalised to the means for mice treated with vehicle only. **D,** Representative images of heart sections stained with picrosirius red. The middle and lower panels illustrate the average and maximum response, respectively, for each condition. **E,** Interstitial fibrosis was measured using Image J and is presented as the % of the total area (excluding regions around the blood vessels). Perivascular fibrosis was scored (1: negligible increase in fibrosis around any vessel; 2: mild to moderate fibrosis around 1 or more vessels; 3: significant fibrosis permeating tissue around 1 or more vessels; 4: extensive fibrosis around multiple vessels, penetrating into the myocardium). **F,** RNA was extracted from mouse heart powders and analysed by qPCR for fibrosis genes (*Col1a1, Col4a1, Fn1* and *Postn1*). Results are relative to GAPDH and normalised to the means for mice treated with vehicle only. Individual datapoints are plotted with means ± SD. Statistical analysis used 2-way ANOVA with Holm-Sidak’s post-test.

## Discussion

Eukaryotic cellular responses are regulated by vast numbers of protein phosphorylation reactions, catalysed by over 500 different protein kinases in the mammalian kinome [43, 44] and countered by a range of protein phosphatases [45]. We have detailed knowledge of how some key signalling pathways operate, but the regulation and roles of many protein kinases remain to be unravelled. Here, we focused on a relatively uninvestigated system, the STRIPAK complexes with a striatin isoform at the core, bringing together the most abundant protein phosphatase in the cell (PP2A) with key protein kinases (GCKs) to regulate their activation [25, 26]. Our data indicate that the three striatin isoforms are all dysregulated in human failing hearts, but our studies with genetically-altered mice place a particular emphasis on striatin itself in the development of cardiac hypertrophy induced by AngII treatment and, therefore, in the broader context of hypertensive heart disease.

The data support a working model in which striatin-based STRIPAKs operate in all cardiac cells (**Figure 7A**). AngII probably acts primarily on endothelial cells since these are highly responsive to AngII and are in direct contact with AngII in the blood. Amongst other effects, AngII stimulates endothelial cells to release pro-hypertrophic factors (e.g. endothelin-1, Edn1 [46]) that act on cardiomyocytes, inducing cardiomyocyte hypertrophy. In turn, cardiomyocytes release factors that promote fibrosis (e.g. fibroblast growth factor 2, FGF2) and proliferation (e.g. EGF family ligands) in other cardiac cells. Our previous studies with BRAF knockout mice support this concept since *Edn1* and *FGF2* mRNAs are upregulated in mouse hearts by AngII, but manipulation of cardiomyocyte signalling (with cardiomyocyte knockout of BRAF) selectively inhibit the increase in *FGF2* [34]. We propose that the response involves striatin-based STRIPAKs with activation of GCKs in one or more of the cardiac cell types. With cardiomyocyte knockout of STRN (**Figure 7B**), the likely dysregulation and mislocalisation of one or more GCK in cardiomyocytes is sufficient to reduce hypertrophic growth (**Figures 5-6**). Pro-fibrotic cardiomyocyte-derived factors are also reduced, resulting in inhibition of fibrotic genes and overall reduction in cardiac fibrosis. With heterozygous global striatin knockout (**Figure 7C**), the situation is more complex. Striatin-based STRIPAKs are decreased in all cardiac cells (**Figure 2**), most likely resulting in some dysregulation and mislocalisation of GCKs throughout the heart. Thus, there is probably reduced production of hypertrophic factors from endothelial cells to suppress cardiomyocyte hypertrophy and, coupled with the effect of striatin loss in cardiomyocytes, this potentially leads to more profound inhibition of cardiomyocyte hypertrophy (**Figure 3**). There is also likely to be a reduction in cardiomyocyte derived pro-fibrotic factors, but we did not detect any decrease in cardiac fibrosis (**Figure 4**). This could be due to dysregulation of the system generally, increasing the overall level of stress on the heart.

**Figure 7.**
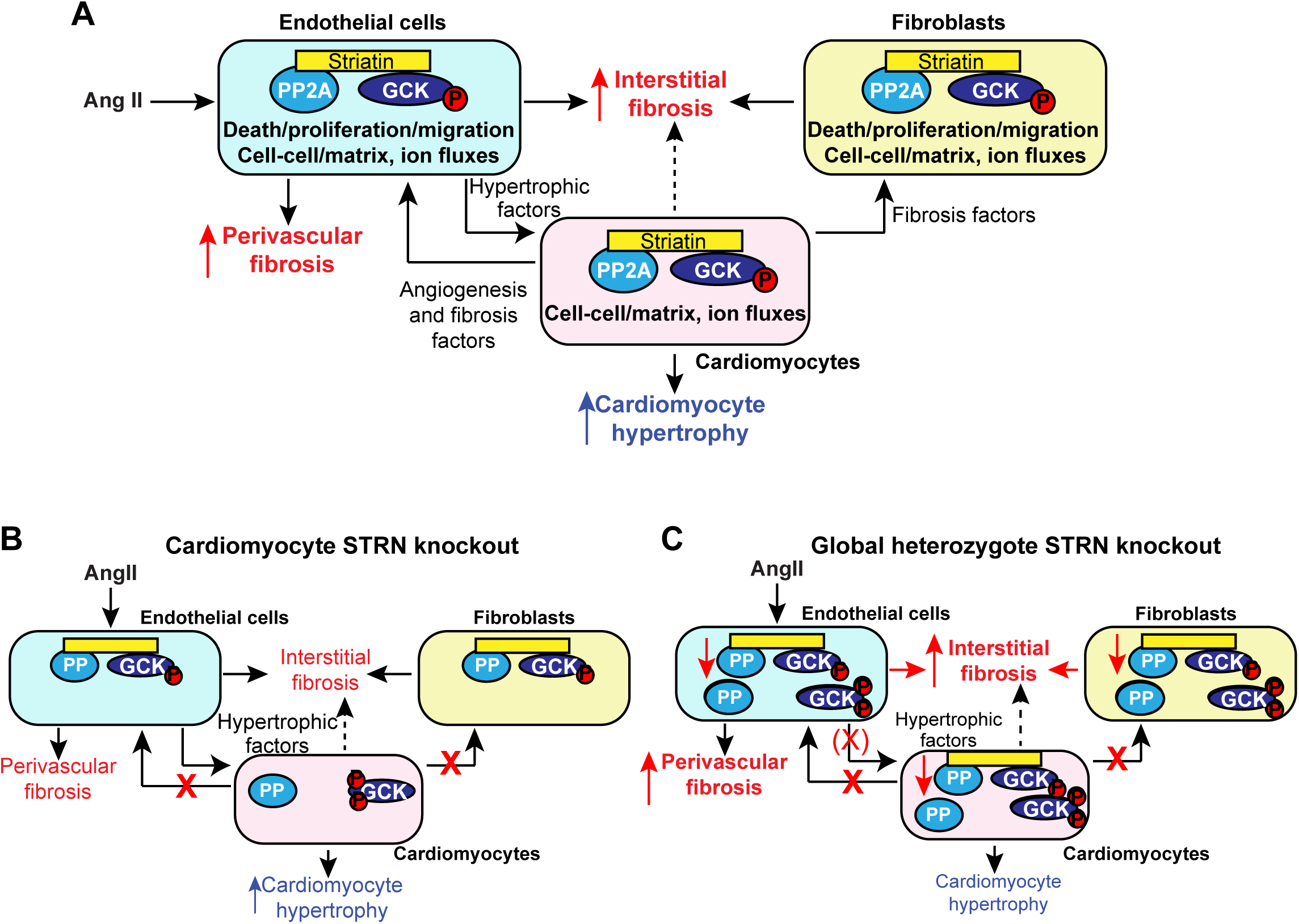
Schematic for the role of striatin in cardiac hypertrophy induced by angiotensin II. Striatin-based STRIPAK complexes link PP2A with protein kinases of the germinal centre kinase (GCK) family. PP2A dephosphorylates GCKs to maintain the inactive state in the three main cell types of the heart (cardiomyocytes, endothelial cells, fibroblasts). GCKs in STRIPAKs are targeted to subcellular domains where they modulate cell-cell and cell-matrix interactions, regulate cell death or (in endothelial cells or fibroblasts) influence cell proliferation and migration. **A**, In our working model, AngII acts primarily on endothelial cells in direct contact with the blood and highly responsive to this hormone. Endothelial cells release pro-hypertrophic factors to promote cardiomyocyte hypertrophy. Cardiomyocytes release additional pro-fibrotic and pro-proliferative factors that affect other cardiac cells. The response involves striatin-based STRIPAKs and activation of GCKs in one or more of the cardiac cell types, with GCKs potentially being activated by reduced PP2A activity in the STRIPAK complex. **B**, Cardiomyocyte-specific knockout of striatin caused dysregulation (with possible increased phosphorylation) and mislocalisation of GCKs in cardiomyocytes, reducing cardiomyocyte hypertrophy. The response was not completely lost because pro-hypertrophic factors acting through other signalling pathways were still produced by cardiac non-myocytes. Cardiomyocyte-derived factors that promote fibrosis were probably reduced, reflected in reduced expression of fibrotic genes and an overall reduction in cardiac fibrosis. **C**, In heterozygous global striatin knockout mice, striatin-based STRIPAKs were decreased in all cardiac cells resulting in some dysregulation and mislocalisation of GCKs. This resulted in reduced production of hypertrophic factors and, coupled with loss of striatin in cardiomyocytes, more profound inhibition of cardiomyocyte hypertrophy. Although there was probably some reduction in cardiomyocyte-derived pro-fibrotic factors, together with dysregulation of STRIPAK components in non-cardiomyocytes, this was not sufficient to suppress cardiac fibrosis overall.

Previous studies in mice with global heterozygous STRN knockout used the same knockout-first system as we used here. The earlier studies developed from prior work demonstrating upregulation of STRN in mouse heart and aorta by aldosterone, modulation of dietary salt or a combination of L-NAME (to inhibit NO production) and AngII [47]. STRN^+/-^ mice and their wild-type littermates have similar blood pressure when fed a restricted salt or normal diet (0.03% or 0.3-0.5% NaCl, respectively), but have an exaggerated increase in blood pressure on a higher salt diet (1.6% NaCl) along with enhanced contraction of aortic rings and reduced relaxation [3–5]. STRN^+/-^ mice also have an enhanced response to aldosterone with increased renal (though not cardiac) damage [5]. Interestingly, there is no significant difference in the renal or cardiac responses of STRN^+/-^ and wild-type mice in a hypertension model using L-NAME and AngII, despite an increase in blood pressure. This apparently contradicts our data, but the models are different. L-NAME specifically inhibits NO production and compromises endothelial cell function and vessel relaxation [48]. In the Garza et al. model [4], mice were treated with L-NAME for 7 d, before implantation of minipumps for delivery of 0.7 mg/kg/d AngII for 3 d in a regime that increased blood pressure. Our approach used AngII at a similar dose over 7 d, but without L-NAME, a regime with a minimal effect on blood pressure over this time [39–41] and, arguably, a milder model. Our goal was to assess effects of AngII as hypertension develops and our data indicate that STRN is required for this early adaptive phase of the cardiac response.

STRN3 was first identified as a nuclear antigen (S/G2 nuclear antigen, SG2NA) subject to cell cycle regulation [49] and, consequently, there is greater emphasis on its role in proliferating cells and cancer (e.g. [50]). However, it is expressed at significant levels in the heart and, of the multiple splice-variants, the dominant isoform is reported to be 78 kD, lacking two exons from the full-length 87 kDa variant [51, 52] Although other variants are reported in the heart, we detected a single dominant band above the 75 kDa marker, presumably corresponding to the 78 kDa STRN3 isoform (**Supplementary Figures S1-S3**). We are not aware of other published studies of STRN3 in the heart or of *in vivo* studies in STRN3 knockout mice. STRN3 was expressed in mouse and human hearts and was significantly upregulated in human failing hearts (**Figure 1A-C**). However, we did not detect any significant differences in cardiac function or dimensions between STRN3^+/-^ mice and their wild-type littermates, either at baseline or in response to AngII (**Figures 2-4**). The studies of STRN3^+/-^ and STRN^+/-^ mice were done in parallel and the negative results with STRN3^+/-^ mice emphasize the potential importance of STRN in the early adaptive response to AngII. However, STRN3 may play an important role in later phases of hypertension-induced cardiac dysfunction and/or in other cardiac pathologies (e.g. myocardial infarction that results in acute injury). It also has to be considered that we studied mice with heterozygote rather than homozygote gene deletion and study of homozygotes (not possible because of embryonic lethality) may have been more revealing. Because of this, it was important to adopt a conditional deletion approach to avoid problems during development.

Immunostaining studies place STRN at the intercalated disc in the heart suggesting it may regulate ion fluxes [6, 30]. Consistent with this concept, reduced expression of STRN in boxer dogs is associated with ARVC and heart failure [6, 7], both of which are associated with a higher risk of life-threatening ventricular arrhythmia and poor prognosis [53]. In addition, studies in cultured cardiomyocytes indicate that overexpression of striatin enhances contraction and STRN knockdown reduces contractility [54]. Human genome-wide association studies (GWAS) link the STRN gene with QRS/PR interval [10, 11], further suggesting a role in regulating ion fluxes and contractility in cardiomyocytes. We did not detect any differences in cardiac function between STRN^+/-^ mice and their wild-type littermates, but it is unlikely that we would have detected arrhythmias with echocardiography in the relatively young mice we studied with a relatively low level of stress resulting from the dose and duration of AngII treatment. Greater differences would perhaps have been detected in older mice or with a more severe or prolonged stress (e.g. in a myocardial infarction model, prolonged treatment with AngII or with transverse aortic constriction).

Even though we saw no effect of heterozygous STRN knockout on cardiac function using echocardiography, the overall hypertrophic response induced by AngII was inhibited (**Figure 2**). This was due to a reduction in cardiomyocyte hypertrophy rather than fibrosis (**Figures 3-4**), suggesting that the phenotype resulted from decreased STRN expression in cardiomyocytes. To address this, we developed a model for homozygous cardiomyocyte-specific STRN knockout adopting a system for inducible and conditional gene deletion. This used a well-established approach using a tamoxifen-inducible Cre enzyme under the regulation of the MYH6 promoter [36], a system which avoids problems associated with development but raises additional concerns of potential cardiotoxicity from the Cre enzyme. This was minimised by only using mice hemizygous for Cre and by restraining temporal activation of the enzyme with just a single dose of tamoxifen to induce recombination. In hemizygous Cre^+/-^ mice, we detect no cardiotoxicity with or without AngII for at least the duration of the experiments reported here [33, 35]. Others have used a similar approach and also report little cardiotoxicity [55]. Given the results with STRN^+/-^ mice, we anticipated that the hypertrophic response induced by AngII would be compromised by cardiomyocyte STRN knockout, and the increase in predicted LV mass estimated on echocardiograms was, indeed, reduced (**Figure 5G**). However, the degree of inhibition of cardiomyocyte hypertrophy appeared less than with the STRN^+/-^ mice and, in contrast to the STRN^+/-^ mice, there was substantial reduction in fibrosis (**Figure 6**). The mice were derived from sperm from our STRN^+/-^ colony so the difference is unlikely to be due to the genetic background. Thus, the effect on fibrosis is most probably a true reflection of the knockout system, leading to our working model in **Figure 7** as described above.

SNPs in the STRN gene have been linked to regulation of blood pressure and the development of heart failure using GWAS, but there are some difficulties with interpretation. The first SNP to be linked to QRS interval (rs17020136 [56]) was originally placed in the STRN gene, but is now linked to the adjacent HEATR5B gene in the EMBL-EBI GWAS Catalog (https://www.ebi.ac.uk/gwas), along with others associated with increased systolic blood pressure (rs146074994, 13408514 [9, 57]). HEATR5B (HEAT repeat containing 5B) is a ubiquitously expressed protein-coding gene of unknown function and further studies of its role in blood pressure regulation may be useful. Nevertheless, SNPs in the STRN gene are also linked to increased blood pressure (rs2540923 [3], rs3770770 [9]) in addition to QRS/PR interval (rs3770770 [10], rs17496249 [11]), hypertrophic cardiomyopathy and heart failure (rs2003585 [12]). Many of the identified STRN SNPs associated with cardiac dysfunction are intronic, so the functional consequences are not clear. Nevertheless, linkage of the STRN gene with blood pressure along with studies in STRN^+/-^ mice have led to a clinical trial for use of mineralocorticoid receptor antagonists in hypertensive patients carrying STRN risk alleles [58].

Our data implicate cardiomyocyte STRN in cardiac hypertrophy, but provide limited insight into the mechanism of action. Striatin itself becomes hyperphosphorylated on inhibition of PP2A in cardiomyocytes [59], a modification which may modify subcellular localisation and/or binding partners. It may also be subject to ADP-ribosylation [60], although this has not been studied in the heart. The protein kinases identified in STRIPAK complex signalling belong to the GCK family with the GCKII (MST1 and MST2 [61]), GCKIII (MST3, MST4, YSK [50, 62–64]), GCKV (SLIK in Drosophila; SLK and LOK are mammalian homologues [63, 65]) and GCKVI (MAP4K4, TNIK, MINK1 [66–68]) subfamilies being specifically implicated to date. MST1/2, MST3, SLK and MAP4K4 are relatively highly expressed in adult rat cardiomyocytes [69], so these are the candidate kinases for cardiac adaptation to AngII. MST1/MST2 are involved in HIPPO signalling and regulation of cell survival/cell death in the heart [70]. Since cardiomyocyte MST1 knockout increases autophagic flux to alleviate AngII-induced cardiac damage [71], dysregulation of MST1 as a result of cardiomyocyte STRN knockout could have a similar effect and reduce cardiomyocyte hypertrophy. MAP4K4 associates with striatins in cardiomyocytes and is linked to human heart failure [59, 72, 73], so could also be involved. MST3 plays an important role in cell migration and is regulated acutely by phosphatase activity in cardiomyocytes [74, 75], but there is little/no information on the role of SLK in heart. Whilst all of the kinases may interact with each of the striatins in experiments conducted *in vitro* or using overexpression approaches, specificity in terms of STRIPAK binding partners or subcellular targeting remains to be determined.

This study only considers the role of STRN and STRN3 in the early stages of cardiac remodelling induced by AngII, not the later stages associated with heart failure and decreased ejection fraction. Extending the study over a more prolonged period would enable further assessment of whether STRN knockdown or cardiomyocyte deletion could prevent this deterioration of cardiac function. Longer term studies would also help to determine if STRN3 plays an important role in developing heart failure, as suggested by the minor abnormalities in longitudinal strain we detected in STRN3^+/-^ mice treated with AngII over 7 d. We also did not consider the possible effects of STRN on arrhythmias and sudden death. We noted that the STRN mice under investigation in this study appeared more prone to sudden death than other genetically altered mice (e.g. those associated with BRAF [33, 35]) we studied in parallel. However, there was no correlation with STRN expression (**Supplementary Table S1**), suggesting it was either coincidental or related to the background strain. Given the link between STRN and ARVC in boxer dogs [6, 7] and SNPs in the STRN gene to hypertrophic cardiomyopathy, it will be important to conduct additional studies to assess possible arrhythmias in mice with STRN knockdown. Probably the greatest limitation of this study is the lack of knowledge of the STRIPAK complexes themselves. Thus, although the data suggest that inhibiting STRN will reduce cardiac hypertrophy induced by AngII, STRN potentially acts at the core of multiple complexes that regulate different GCKs, and one or more of these GCKs may be involved. Knowing which GCKs are involved and how they are regulated will be a crucial element for identifying specific targets for therapeutic manipulation of STRIPAK signalling.

In conclusion, our data indicate that STRN, but probably not STRN3, plays an important role in the early remodelling processes induced in the heart to AngII. Although there is clearly much research still to be done, the data clearly identify the striatin-based STRIPAK system as a novel signalling paradigm in development of pathological cardiac hypertrophy. Understanding this system may provide therapeutic options for modulating the responses and managing progression of hypertensive heart disease.

## Funding

This work was supported by the British Heart Foundation (PG/15/41/31560, FS/18/33/33621, PG/15/24/31367, FS/19/24/34262) and Qassim University, Saudi Arabia (to H.O.A).

## Acknowledgements

We thank Andrew Cripps, Mhairi Baxter and Wayne Knight (University of Reading), and Robert Bond, Emma Mustafa and Rene Ocho (St. George’s University of London) for support for the *in vivo* mouse studies.

## Author contributions

J.J.C. was responsible for and conducted the breeding for the experiments with the STRN^+/-^ and STRN3^+/-^ mice, and the echocardiography and analysis. H.O.A. assisted with these experiments. Studies with mice for cardiomyocyte knockout of STRN were conducted by J.J.C and S.T.E.C. S.P.C and O.J.L.R were responsible for the analysis of RNASeq data from dilated cardiomyopathy patients compared with normal controls. P.H.S., N.R, R.K and P.R.D. assisted with writing and reviewing the manuscript. P.H.S. and A.C. initiated the studies. A.C. obtained funding, designed the experiments and wrote the manuscript.

## Data Availability Statement

All primary data are available from the corresponding author upon reasonable request.

## Conflict of interests

The authors declare no conflicts of interests.

## Notes

### Competing Interest Statement

The authors have declared no competing interest.

## References

1 Savarese, G., Becher, P.M., Lund, L.H., Seferovic, P., Rosano, G.M.C. and Coats, A. (2023) Global burden of heart failure: A comprehensive and updated review of epidemiology. Cardiovasc Res. 118, 3272–3287, 10.1093/cvr/cvac013

2 Mills, K.T., Stefanescu, A. and He, J. (2020) The global epidemiology of hypertension. Nat Rev Nephrol. 16, 223–237, 10.1038/s41581-019-0244-2

3 Garza, A.E., Rariy, C.M., Sun, B., Williams, J.S., Lasky-Su, J., Baudrand, R. et al. (2015) Variants in striatin gene are associated with salt-sensitive blood pressure in mice and humans. Hypertension. 65, 211–217, 10.1161/HYPERTENSIONAHA.114.04233

4 Garza, A.E., Pojoga, L.H., Moize, B., Hafiz, W.M., Opsasnick, L.A., Siddiqui, W.T. et al. (2015) Critical role of striatin in blood pressure and vascular responses to dietary sodium intake. Hypertension. 66, 674–680, 10.1161/HYPERTENSIONAHA.115.05600

5 Garza, A.E., Trefts, E., Katayama Rangel, I.A., Brooks, D., Baudrand, R., Moize, B. et al. (2020) Striatin heterozygous mice are more sensitive to aldosterone-induced injury. J Endocrinol. 245, 439–450, 10.1530/JOE-19-0562

6 Meurs, K.M., Mauceli, E., Lahmers, S., Acland, G.M., White, S.N. and Lindblad-Toh, K. (2010) Genome-wide association identifies a deletion in the 3’ untranslated region of striatin in a canine model of arrhythmogenic right ventricular cardiomyopathy. Hum Genet. 128, 315–324, 10.1007/s00439-010-0855-y

7 Meurs, K.M., Stern, J.A., Sisson, D.D., Kittleson, M.D., Cunningham, S.M., Ames, M.K. et al. (2013) Association of dilated cardiomyopathy with the striatin mutation genotype in boxer dogs. J Vet Intern Med. 27, 1437–1440, 10.1111/jvim.12163

8 Gupta, T., Connors, M., Tan, J.W., Manosroi, W., Ahmed, N., Ting, P.Y. et al. (2017) Striatin gene polymorphic variants are associated with salt sensitive blood pressure in normotensives and hypertensives. Am J Hypertens. 31, 124–131, 10.1093/ajh/hpx146

9 Plotnikov, D., Huang, Y., Khawaja, A.P., Foster, P.J., Zhu, Z., Guggenheim, J.A. et al. (2022) High blood pressure and intraocular pressure: a Mendelian randomization study. Invest Ophthalmol Vis Sci. 63, 29, 10.1167/iovs.63.6.29

10 van der Harst, P., van Setten, J., Verweij, N., Vogler, G., Franke, L., Maurano, M.T., et al. (2016) 52 Genetic loci influencing myocardial mass. J Am Coll Cardiol. 68, 1435–1448, 10.1016/j.jacc.2016.07.729

11 Ntalla, I., Weng, L.C., Cartwright, J.H., Hall, A.W., Sveinbjornsson, G., Tucker, N.R. et al. (2020) Multi-ancestry GWAS of the electrocardiographic PR interval identifies 202 loci underlying cardiac conduction. Nat Commun. 11, 2542, 10.1038/s41467-020-15706-x

12 Harper, A.R., Goel, A., Grace, C., Thomson, K.L., Petersen, S.E., Xu, X. et al. (2021) Common genetic variants and modifiable risk factors underpin hypertrophic cardiomyopathy susceptibility and expressivity. Nat Genet. 53, 135–142, 10.1038/s41588-020-00764-0

13 Levin, M.G., Tsao, N.L., Singhal, P., Liu, C., Vy, H.M.T., Paranjpe, I. et al. (2022) Genome-wide association and multi-trait analyses characterize the common genetic architecture of heart failure. Nat Commun. 13, 6914, 10.1038/s41467-022-34216-6

14 Surendran, P., Feofanova, E.V., Lahrouchi, N., Ntalla, I., Karthikeyan, S., Cook, J. et al. (2020) Discovery of rare variants associated with blood pressure regulation through meta-analysis of 1.3 million individuals. Nat Genet. 52, 1314–1332, 10.1038/s41588-020-00713-x

15 Zhou, P. and Pu, W.T. (2016) Recounting cardiac cellular composition. Circ Res. 118, 368–370, 10.1161/CIRCRESAHA.116.308139

16 Dorn, G.W., II, Robbins, J. and Sugden, P.H. (2003) Phenotyping hypertrophy: eschew obfuscation. Circ Res. 92, 1171–1175, 10.1161/01.RES.0000077012.11088.BC

17 Sheng, S.Y., Li, J.M., Hu, X.Y. and Wang, Y. (2023) Regulated cell death pathways in cardiomyopathy. Acta Pharmacol Sin. 44, 1521–1535, 10.1038/s41401-023-01068-9

18 Gogiraju, R., Bochenek, M.L. and Schafer, K. (2019) Angiogenic endothelial cell signaling in cardiac hypertrophy and heart failure. Front Cardiovasc Med. 6, 20, 10.3389/fcvm.2019.00020

19 Suthahar, N., Meijers, W.C., Sillje, H.H.W. and de Boer, R.A. (2017) From inflammation to fibrosis-molecular and cellular mechanisms of myocardial tissue remodelling and perspectives on differential treatment opportunities. Curr Heart Fail Rep. 14, 235–250, 10.1007/s11897-017-0343-y

20 Mishra, S. and Kass, D.A. (2021) Cellular and molecular pathobiology of heart failure with preserved ejection fraction. Nat Rev Cardiol. 18, 400–423, 10.1038/s41569-020-00480-6

21 Kurose, H. (2021) Cardiac fibrosis and fibroblasts. Cells. 10, 1716, 10.3390/cells10071716

22 Kovacic, J.C., Dimmeler, S., Harvey, R.P., Finkel, T., Aikawa, E., Krenning, G. et al. (2019) Endothelial to mesenchymal transition in cardiovascular disease: JACC State-of-the-Art Review. J Am Coll Cardiol. 73, 190–209, 10.1016/j.jacc.2018.09.089

23 Reynhout, S. and Janssens, V. (2019) Physiologic functions of PP2A: Lessons from genetically modified mice. Biochim Biophys Acta Mol Cell Res. 1866, 31–50, 10.1016/j.bbamcr.2018.07.010

24 Delpire, E. (2009) The mammalian family of sterile 20p-like protein kinases. Pflugers Archives. 458, 953–967, 10.1007/s00424-009-0674-y

25 Hwang, J. and Pallas, D.C. (2014) STRIPAK complexes: structure, biological function, and involvement in human diseases. Int J Biochem Cell Biol. 47, 118–148, 10.1016/j.biocel.2013.11.021

26 Kuck, U., Radchenko, D. and Teichert, I. (2019) STRIPAK, a highly conserved signaling complex, controls multiple eukaryotic cellular and developmental processes and is linked with human diseases. Biol Chem. 400, 1005–1022, 10.1515/hsz-2019-0173

27 Goudreault, M., D’Ambrosio, L.M., Kean, M.J., Mullin, M.J., Larsen, B.G., Sanchez, A. et al. (2009) A PP2A phosphatase high density interaction network identifies a novel striatin-interacting phosphatase and kinase complex linked to the cerebral cavernous malformation 3 (CCM3) protein. Mol Cell Proteomics. 8, 157–171, 10.1074/mcp.M800266-MCP200

28 Herzog, F., Kahraman, A., Boehringer, D., Mak, R., Bracher, A., Walzthoeni, T. et al. (2012) Structural probing of a protein phosphatase 2A network by chemical cross-linking and mass spectrometry. Science. 337, 1348–1352, 10.1126/science.1221483

29 Couzens, A.L., Knight, J.D., Kean, M.J., Teo, G., Weiss, A., Dunham, W.H. et al. (2013) Protein interaction network of the mammalian Hippo pathway reveals mechanisms of kinase-phosphatase interactions. Sci Signal. 6, rs15, 10.1126/scisignal.2004712

30 Franke, W.W., Rickelt, S., Zimbelmann, R., Dorflinger, Y., Kuhn, C., Frey, N. et al. (2014) Striatins as plaque molecules of zonulae adhaerentes in simple epithelia, of tessellate junctions in stratified epithelia, of cardiac composite junctions and of various size classes of lateral adherens junctions in cultures of epithelia- and carcinoma-derived cells. Cell Tissue Res. 357, 645–665, 10.1007/s00441-014-2053-z

31 Heinig, M., Adriaens, M.E., Schafer, S., van Deutekom, H.W.M., Lodder, E.M., Ware, J.S., et al. (2017) Natural genetic variation of the cardiac transcriptome in non-diseased donors and patients with dilated cardiomyopathy. Genome Biol. 18, 170, 10.1186/s13059-017-1286-z

32 Love, M.I., Huber, W. and Anders, S. (2014) Moderated estimation of fold change and dispersion for RNA-seq data with DESeq2. Genome Biol. 15, 550, 10.1186/s13059-014-0550-8

33 Clerk, A., Meijles, D.N., Hardyman, M.A., Fuller, S.J., Chothani, S.P., Cull, J.J. et al. (2022) Cardiomyocyte BRAF and type 1 RAF inhibitors promote cardiomyocyte and cardiac hypertrophy in mice in vivo. Biochem J. 479, 401–424, 10.1042/BCJ20210615

34 Marshall, J.J., Cull, J.J., Alharbi, H.O., Zaw Thin, M., Cooper, S.T., Barrington, C. et al. (2022) PKN2 deficiency leads both to prenatal congenital cardiomyopathy and defective angiotensin II stress responses. Biochem J. 479, 1467–1486, 10.1042/BCJ20220281

35 Alharbi, H.O., Hardyman, M.A., Cull, J.J., Markou, T., Cooper, S.T.E., Glennon, P.E. et al. (2022) Cardiomyocyte BRAF is a key signalling intermediate in cardiac hypertrophy in mice. Clin Sci (Lond). 136, 1661–1681, 10.1042/CS20220607

36 Sohal, D.S., Nghiem, M., Crackower, M.A., Witt, S.A., Kimball, T.R., Tymitz, K.M. et al. (2001) Temporally regulated and tissue-specific gene manipulations in the adult and embryonic heart using a tamoxifen-inducible Cre protein. Circ Res. 89, 20–25, 10.1161/hh1301.092687

37 Meijles, D.N., Cull, J.J., Markou, T., Cooper, S.T.E., Haines, Z.H.R., Fuller, S.J. et al. (2020) Redox regulation of cardiac ASK1 (Apoptosis Signal-Regulating Kinase 1) controls p38-MAPK (mitogen-activated protein kinase) and orchestrates cardiac remodeling to hypertension. Hypertension. 76, 1208–1218, 10.1161/HYPERTENSIONAHA.119.14556

38 Marshall, A.K., Barrett, O.P.T., Cullingford, T.E., Shanmugasundram, A., Sugden, P.H. and Clerk, A. (2010) ERK1/2 signaling dominates over RhoA signaling in regulating early changes in RNA expression induced by endothelin-1 in neonatal rat cardiomyocytes. PLoS One. 5, e10027, 10.1371/journal.pone.0010027

39 Zimmerman, M.C., Lazartigues, E., Sharma, R.V. and Davisson, R.L. (2004) Hypertension caused by angiotensin II infusion involves increased superoxide production in the central nervous system. Circ Res. 95, 210–216, 10.1161/01.RES.0000135483.12297.e4

40 Patel, J., Douglas, G., Kerr, A.G., Hale, A.B. and Channon, K.M. (2018) Effect of irradiation and bone marrow transplantation on angiotensin II-induced aortic inflammation in ApoE knockout mice. Atherosclerosis. 276, 74–82, 10.1016/j.atherosclerosis.2018.07.019

41 Capone, C., Faraco, G., Peterson, J.R., Coleman, C., Anrather, J., Milner, T.A. et al. (2012) Central cardiovascular circuits contribute to the neurovascular dysfunction in angiotensin II hypertension. J Neurosci. 32, 4878–4886, 10.1523/JNEUROSCI.6262-11.2012

42 Jahn, H.M., Kasakow, C.V., Helfer, A., Michely, J., Verkhratsky, A., Maurer, H.H. et al. (2018) Refined protocols of tamoxifen injection for inducible DNA recombination in mouse astroglia. Sci Rep. 8, 5913, 10.1038/s41598-018-24085-9

43 Manning, G., Whyte, D.B., Martinez, R., Hunter, T. and Sudarsanam, S. (2002) The protein kinase complement of the human genome. Science. 298, 1912–1934, 10.1126/science.1075762

44 Caenepeel, S., Charydczak, G., Sudarsanam, S., Hunter, T. and Manning, G. (2004) The mouse kinome: discovery and comparative genomics of all mouse protein kinases. Proc Natl Acad Sci U.S.A. 101, 11707–11712, 10.1073/pnas.0306880101

45 Nguyen, H. and Kettenbach, A.N. (2023) Substrate and phosphorylation site selection by phosphoprotein phosphatases. Trends Biochem Sci. 48, 713–725, 10.1016/j.tibs.2023.04.004

46 Marasciulo, F.L., Montagnani, M. and Potenza, M.A. (2006) Endothelin-1: the yin and yang on vascular function. Curr Med Chem. 13, 1655–1665, 10.2174/092986706777441968

47 Pojoga, L.H., Coutinho, P., Rivera, A., Yao, T.M., Maldonado, E.R., Youte, R. et al. (2012) Activation of the mineralocorticoid receptor increases striatin levels. Am J Hypertens. 25, 243–249, 10.1038/ajh.2011.197

48 Evora, P.R., Evora, P.M., Celotto, A.C., Rodrigues, A.J. and Joviliano, E.E. (2012) Cardiovascular therapeutics targets on the NO-sGC-cGMP signaling pathway: a critical overview. Curr Drug Targets. 13, 1207–1214, 10.2174/138945012802002348

49 Muro, Y., Chan, E.K., Landberg, G. and Tan, E.M. (1995) A cell-cycle nuclear autoantigen containing WD-40 motifs expressed mainly in S and G2 phase cells. Biochem Biophys Res Commun. 207, 1029–1037, 10.1006/bbrc.1995.1288

50 Madsen, C.D., Hooper, S., Tozluoglu, M., Bruckbauer, A., Fletcher, G., Erler, J.T. et al. (2015) STRIPAK components determine mode of cancer cell migration and metastasis. Nat Cell Biol. 17, 68–80, 10.1038/ncb3083

51 Jain, B.P., Chauhan, P., Tanti, G.K., Singarapu, N., Ghaskadbi, S. and Goswami, S.K. (2015) Tissue specific expression of SG2NA is regulated by differential splicing, RNA editing and differential polyadenylation. Gene. 556, 119–126, 10.1016/j.gene.2014.11.045

52 Sanghamitra, M., Talukder, I., Singarapu, N., Sindhu, K.V., Kateriya, S. and Goswami, S.K. (2008) WD-40 repeat protein SG2NA has multiple splice variants with tissue restricted and growth responsive properties. Gene. 420, 48–56, 10.1016/j.gene.2008.04.016

53 Krahn, A.D., Wilde, A.A.M., Calkins, H., La Gerche, A., Cadrin-Tourigny, J., Roberts, J.D., et al. (2022) Arrhythmogenic Right Ventricular Cardiomyopathy. JACC Clin Electrophysiol. 8, 533–553, 10.1016/j.jacep.2021.12.002

54 Nader, M., Alotaibi, S., Alsolme, E., Khalil, B., Abu-Zaid, A., Alsomali, R. et al. (2017) Cardiac striatin interacts with caveolin-3 and calmodulin in a calcium sensitive manner and regulates cardiomyocyte spontaneous contraction rate. Can J Physiol Pharmacol. 95, 1306–1312, 10.1139/cjpp-2017-0155

55 Hougen, K., Aronsen, J.M., Stokke, M.K., Enger, U., Nygard, S., Andersson, K.B. et al. (2010) Cre-loxP DNA recombination is possible with only minimal unspecific transcriptional changes and without cardiomyopathy in Tg(alphaMHC-MerCreMer) mice. Am J Physiol Heart Circ Physiol. 299, H1671–1678, 10.1152/ajpheart.01155.2009

56 Sotoodehnia, N., Isaacs, A., de Bakker, P.I., Dorr, M., Newton-Cheh, C., Nolte, I.M., et al. (2010) Common variants in 22 loci are associated with QRS duration and cardiac ventricular conduction. Nat.Genet. 42, 1068–1076, 10.1038/ng.716

57 Giri, A., Hellwege, J.N., Keaton, J.M., Park, J., Qiu, C., Warren, H.R. et al. (2019) Trans-ethnic association study of blood pressure determinants in over 750,000 individuals. Nat Genet. 51, 51–62, 10.1038/s41588-018-0303-9

58 Stone, I.B., Green, J., Koefoed, A.W., Hornik, E.S., Williams, J.S., Adler, G.K. et al. (2021) Striatin genotype-based, mineralocorticoid receptor antagonist-driven clinical trial: study rationale and design. Pharmacogenet Genomics. 31, 83–88, 10.1097/FPC.0000000000000425

59 Fuller, S.J., Edmunds, N.S., McGuffin, L.J., Hardyman, M.A., Cull, J.J., Alharbi, H.O. et al. (2021) MAP4K4 expression in cardiomyocytes: multiple isoforms, multiple phosphorylations and interactions with striatins. Biochem J. 478, 2121–2143, 10.1042/BCJ20210003

60 Guettler, S., LaRose, J., Petsalaki, E., Gish, G., Scotter, A., Pawson, T. et al. (2011) Structural basis and sequence rules for substrate recognition by Tankyrase explain the basis for cherubism disease. Cell. 147, 1340–1354, 10.1016/j.cell.2011.10.046

61 Tang, Y., Chen, M., Zhou, L., Ma, J., Li, Y., Zhang, H. et al. (2019) Architecture, substructures, and dynamic assembly of STRIPAK complexes in Hippo signaling. Cell Discov. 5, 3, 10.1038/s41421-018-0077-3

62 Gordon, J., Hwang, J., Carrier, K.J., Jones, C.A., Kern, Q.L., Moreno, C.S. et al. (2011) Protein phosphatase 2a (PP2A) binds within the oligomerization domain of striatin and regulates the phosphorylation and activation of the mammalian Ste20-Like kinase Mst3. BMC Biochem. 12, 54, 10.1186/1471-2091-12-54

63 Kean, M.J., Ceccarelli, D.F., Goudreault, M., Sanches, M., Tate, S., Larsen, B. et al. (2011) Structure-function analysis of core STRIPAK proteins: a signaling complex implicated in Golgi polarization. J Biol Chem. 286, 25065–25075, 10.1074/jbc.M110.214486

64 Ceccarelli, D.F., Laister, R.C., Mulligan, V.K., Kean, M.J., Goudreault, M., Scott, I.C. et al. (2011) CCM3/PDCD10 heterodimerizes with germinal center kinase III (GCKIII) proteins using a mechanism analogous to CCM3 homodimerization. J Biol Chem. 286, 25056–25064, 10.1074/jbc.M110.213777

65 De Jamblinne, C.V., Decelle, B., Dehghani, M., Joseph, M., Sriskandarajah, N., Leguay, K., et al. (2020) STRIPAK regulates Slik localization to control mitotic morphogenesis and epithelial integrity. J Cell Biol. 219, e201911035, 10.1083/jcb.201911035

66 Hyodo, T., Ito, S., Hasegawa, H., Asano, E., Maeda, M., Urano, T. et al. (2012) Misshapen-like kinase 1 (MINK1) is a novel component of striatin-interacting phosphatase and kinase (STRIPAK) and is required for the completion of cytokinesis. J Biol Chem. 287, 25019–25029, 10.1074/jbc.M112.372342

67 Kim, J.W., Berrios, C., Kim, M., Schade, A.E., Adelmant, G., Yeerna, H. et al. (2020) STRIPAK directs PP2A activity toward MAP4K4 to promote oncogenic transformation of human cells. Elife. 9, e53003, 10.7554/eLife.53003

68 Migliavacca, J., Zullig, B., Capdeville, C., Grotzer, M.A. and Baumgartner, M. (2022) Cooperation of striatin 3 and MAP4K4 promotes growth and tissue invasion. Commun Biol. 5, 795, 10.1038/s42003-022-03708-y

69 Fuller, S.J., Osborne, S.A., Leonard, S.J., Hardyman, M.A., Vaniotis, G., Allen, B.G. et al. (2015) Cardiac protein kinases: the cardiomyocyte kinome and differential kinase expression in human failing hearts. Cardiovasc Res. 108, 87–98, 10.1093/cvr/cvv210

70 Wang, J., Liu, S., Heallen, T. and Martin, J.F. (2018) The Hippo pathway in the heart: pivotal roles in development, disease, and regeneration. Nat Rev Cardiol. 15, 672–684, 10.1038/s41569-018-0063-3

71 Cheng, Z., Zhang, M., Hu, J., Lin, J., Feng, X., Wang, S. et al. (2018) Mst1 knockout enhances cardiomyocyte autophagic flux to alleviate angiotensin II-induced cardiac injury independent of angiotensin II receptors. J Mol Cell Cardiol. 125, 117–128, 10.1016/j.yjmcc.2018.08.028

72 Golforoush, P.A., Narasimhan, P., Chaves-Guerrero, P.P., Lawrence, E., Newton, G., Yan, R. et al. (2020) Selective protection of human cardiomyocytes from anthracycline cardiotoxicity by small molecule inhibitors of MAP4K4. Sci Rep. 10, 12060, 10.1038/s41598-020-68907-1

73 Fiedler, L.R., Chapman, K., Xie, M., Maifoshie, E., Jenkins, M., Golforoush, P.A. et al. (2019) MAP4K4 inhibition promotes survival of human stem cell-derived cardiomyocytes and reduces infarct size in vivo. Cell Stem Cell. 24, 579–591 e512, 10.1016/j.stem.2019.01.013

74 Fuller, S.J., McGuffin, L.J., Marshall, A.K., Giraldo, A., Pikkarainen, S., Clerk, A. et al. (2012) A novel non-canonical mechanism of regulation of MST3 (mammalian Sterile20-related kinase 3). Biochem J. 442, 595–610, 10.1042/BJ20112000

75 Sugden, P.H., McGuffin, L.J. and Clerk, A. (2013) SOcK, MiSTs, MASK and STicKs: the germinal centre kinase III (GCKIII) kinases and their heterologous protein-protein interactions. Biochem J. 454, 13–30, 10.1042/BJ20130219

