## Supplemental information for "Striatin plays a major role in angiotensin II-induced cardiomyocyte and cardiac hypertrophy in mice *in vivo*"

### STRN, but not STRN3, plays a major role in angiotensin II-induced cardiomyocyte and cardiac hypertrophy

**Supplementary Table S1. Mouse weights.**

**Supplementary Table S2. Primers for genotyping and confirmation of recombination.**

**Supplementary S3. qPCR primers.**

**Supplementary Table S4. Baseline echocardiography data for $STRN^{+/-}$ and $STRN3^{+/-}$ male mice and their wild-type ($STRN^{+/+}$ and $STRN3^{+/+}$) male littermates.**

**Supplementary Table S5. Echocardiography data for $STRN^{+/-}$ and $STRN3^{+/-}$ male mice and wild-type ($STRN^{+/+}$ and $STRN3^{+/+}$) male littermates treated with acidified PBS (AcPBS) or 0.8 mg/kg/d AngII for 7 d.**

**Supplementary Table S6. Echocardiography data for $STRN3^{fl/fl}/Cre^{+/-}$ mice.**

**Supplementary Figure S1. Full images for immunoblots in Figure 1.**

**Supplementary Figure S2. Full images for immunoblots in Figure 2.**

**Supplementary Figure S3. Full images for immunoblots in Figure 5.**

**Supplementary Table S1. Mouse weights.** Male mice (8 weeks) were allocated to groups on a random basis. $STRN^{+/-}$ and $STRN3^{+/-}$ heterozygous knockout mice and their wild-type (WT) $STRN^{+/+}$ and $STRN3^{+/+}$ littermates were treated with acidified PBS (AcPBS) or 0.8 mg/kg/d angiotensin II (AngII). $STRN^{fl/fl}/Cre^{+/-}$ mice were treated with 40 mg/kg tamoxifen in corn-oil or with corn-oil vehicle (COV) with or without AcPBS or AngII. Weights (g) were taken at the start of the study with the first baseline echocardiogram (Start), immediately after minipump surgery, and when mice were culled (End). Weights post-surgery and at the end included the minipumps. N values are provided with exclusions due to mortality indicated in parentheses

| **Study** | **Condition** | **Start** | | **Post-minipump** | | **End** | | |
|---|---|---|---|---|---|---|---|---|
| | | **Mean** | **SD** | **Mean** | **SD** | **Mean** | **SD** | **n** |
| **STRN study group** | | | | | | | | |
| $STRN^{+/+}$ (WT) | AcPBS | 25.16 | 1.38 | 26.94 | 1.46 | 27.59 | 1.49 | 16 |
| | AngII | 24.77 | 1.57 | 26.46 | 1.42 | 26.65 | 1.38 | 15 (1) |
| $STRN^{+/-}$ | AcPBS | 25.00 | 0.92 | 26.76 | 0.95 | 27.52 | 0.78 | 10 |
| | AngII | 24.78 | 1.38 | 26.49 | 1.26 | 26.85 | 1.57 | 17 |
| **STRN3 study group** | | | | | | | | |
| $STRN3^{+/+}$ (WT) | AcPBS | 23.62 | 2.11 | 25.44 | 1.71 | 26.07 | 1.66 | 16 |
| | AngII | 24.31 | 1.46 | 25.79 | 1.44 | 26.41 | 1.69 | 14 |
| $STRN3^{+/-}$ | AcPBS | 23.26 | 2.20 | 24.85 | 2.01 | 25.76 | 2.15 | 11 |
| | AngII | 23.32 | 1.68 | 25.64 | 1.43 | 25.78 | 1.21 | 11 |
| **$STRN^{fl/fl}/Cre^{+/-}$** | COV/AcPBS | 22.21 | 1.53 | 24.58 | 2.06 | 25.27 | 1.76 | 10 |
| | Tx/AcPBS | 22.82 | 1.70 | 25.79 | 1.73 | 26.40 | 1.31 | 10 |
| | COV/AngII | 21.98 | 2.04 | 24.12 | 1.61 | 24.04 | 1.66 | 9 (2) |
| | Tx/AngII | 25.38 | 1.70 | 25.38 | 1.70 | 25.48 | 1.86 | 8 (1) |

**Supplementary Table S2. Primers for genotyping and confirmation of recombination.**

| **Mouse strain** | **Forward primer** | **Reverse primer** | **Size (bp)** | **Annealing temp.** |
|---|---|---|---|---|
| **Genotyping** | | | | |
| STRN knockout | GAGATGGCGCAACGCAATTAATG | GAAGTGCATGGGAAGTCAGTACACG | 296 | 51°C |
| STRN3 knockout | GAGATGGCGCAACGCAATTAATG | ACCTGAGCCAAATTCACCCAAAACC | 334 | 51°C |
| $Cre^{-}$ | TCTATTGCACACAGCAATCCA | CCAACTCTTGTGAGAGGAGCA | 305 | 52°C |
| $Cre^{+}$ | TCTATTGCACACAGCAATCCA | CCAGCATTGTGAGAACAAGG | 285 | 52°C |
| **Recombination** | | | | |
| Post-FLP | TGAATTATTGGAGTTTTGTTTCAGACC | GCACAGACAGACCTTCATGCTAACC | 630 | 53°C |
| Post-FLP Post-Cre | TGAATTATTGGAGTTTTGTTTCAGACC | GAAGTGCATGGGAAGTCAGTACACG | 666 | 53°C |

#### Supplementary S3. qPCR primers.

| Gene Symbol | Accession No. | Sense Primer (5'→3') | Antisense Primer (5'→3') |
|---|---|---|---|
| Col1a1 | NM_007742 | TCGTGGCTTCTCTGGTCTC | CCGTTGAGTCCGTCTTTGC |
| Col4a1 | NM_009931.2 | TGTGGGCCAGCCAGGCATTG | CAGGGGGTCCGATCGCTCCA |
| Fn1 | NM_010233 | AAGAGGACGTTGCAGAGCTA | AGACACTGGAGACACTGACTAA |
| Gapdh | NM_008084.2 | TCACCACCATGGAGAAGGC | GCTAAGCAGTTGGTGGTGCA |
| Myh7 | NM_080728 | CATGCCAACCGTATGGCTG | GTTCCACGATGGCGATGTTC |
| Nppb | NM_008726 | TCCAGCAGAGACCTCAAAATTC | CAGTGCGTTACAGCCCAAA |
| Postn | NM_015784 | TTCCTCTCCTGCCCTTATATGC | CCTGATCCCGACCCCTGAT |

**Supplementary Table S4. Baseline echocardiography data for $STRN^{+/-}$ and $STRN3^{+/-}$ male mice and their wild-type ($STRN^{+/+}$ and $STRN3^{+/+}$) male littermates.** Long axis B-mode images were analysed using speckle-tracking software. EDV, End diastolic volume; ESV, End systolic volume; EDLVM, End diastolic left ventricle mass; ESLVM, End systolic left ventricle mass; GLS, global longitudinal strain; GCS, Global circumferential strain.

| | $STRN^{+/+}$ (n=31) | | $STRN^{+/-}$ (n=26) | | $STRN3^{+/+}$ (n=27) | | $STRN3^{+/-}$ (n=19) | |
|---|---|---|---|---|---|---|---|---|
| | Mean | SD | Mean | SD | Mean | SD | Mean | SD |
| Heart Rate (bpm) | 484 | 31 | 473 | 54 | 480 | 40 | 472 | 40 |
| Stroke volume (µl) | 29.96 | 5.84 | 31.49 | 6.06 | 28.89 | 4.64 | 28.16 | 7.25 |
| Cardiac output (ml/min) | 14.46 | 2.85 | 14.99 | 3.82 | 13.87 | 2.47 | 13.13 | 2.71 |
| Ejection fraction (%) | 57.54 | 6.86 | 58.14 | 6.45 | 56.26 | 8.10 | 57.38 | 7.83 |
| Fractional shortening (%) | 30.73 | 5.68 | 30.71 | 4.96 | 28.34 | 6.38 | 30.24 | 5.82 |
| EDV (µl) | 52.07 | 9.17 | 54.17 | 9.96 | 51.57 | 8.40 | 48.86 | 9.88 |
| ESV (µl) | 22.12 | 5.61 | 22.68 | 5.88 | 22.68 | 6.52 | 20.70 | 5.43 |
| EDLVM (mg) | 53.90 | 5.43 | 54.21 | 5.83 | 53.24 | 5.20 | 50.75 | 5.79 |
| ESLVM (mg) | 56.03 | 5.76 | 56.11 | 6.20 | 55.99 | 5.94 | 52.99 | 6.01 |
| GLS (%) | -20.27 | 2.91 | -19.80 | 3.44 | -20.76 | 4.97 | -20.06 | 3.16 |
| GCS (%) | -20.84 | 3.48 | -20.83 | 3.40 | -20.30 | 4.01 | -19.98 | 3.34 |

**Supplementary Table S5. Echocardiography data for STRN$^{+/-}$ and STRN3$^{+/-}$ male mice and wild-type (STRN$^{+/+}$ and STRN3$^{+/+}$) male littermates treated with acidified PBS (AcPBS) or 0.8 mg/kg/d AngII for 7 d.** Long axis B-mode images were analysed using speckle-tracking software. EDV, End diastolic volume; ESV, End systolic volume; EDLVM, End diastolic left ventricle mass; ESLVM, End systolic left ventricle mass; GLS, global longitudinal strain; GCS, Global circumferential strain.

| | **STRN$^{+/+}$/AcPBS (n=16)** | | **STRN$^{+/-}$/AcPBS (n=10)** | | **STRN$^{+/+}$/AngII (n=15)** | | **STRN$^{+/-}$/AngII (n=16)** | |
|---|---|---|---|---|---|---|---|---|
| | Mean | SD | Mean | SD | Mean | SD | Mean | SD |
| Heart Rate (bpm) | 507 | 35 | 502 | 38 | 541 | 45 | 526 | 49 |
| Stroke volume (μl) | 29.23 | 5.17 | 28.53 | 5.11 | 25.68 | 5.62 | 27.55 | 5.01 |
| Cardiac output (ml/min) | 14.81 | 2.75 | 14.34 | 2.93 | 13.82 | 2.95 | 14.44 | 2.77 |
| Ejection fraction (%) | 58.93 | 7.05 | 58.94 | 5.63 | 58.19 | 6.84 | 61.99 | 6.51 |
| Fractional shortening (%) | 31.68 | 7.28 | 30.46 | 5.10 | 30.96 | 4.65 | 33.88 | 5.27 |
| EDV (μl) | 49.58 | 8.16 | 48.05 | 6.21 | 44.21 | 9.66 | 44.80 | 9.51 |
| ESV (μl) | 20.35 | 5.27 | 19.52 | 3.24 | 18.53 | 5.82 | 17.25 | 5.49 |
| EDLVM (mg) | 57.23 | 5.12 | 54.27 | 4.70 | 69.82 | 6.81 | 63.32 | 7.95 |
| ESLVM (mg) | 59.03 | 5.17 | 55.47 | 4.94 | 72.29 | 7.61 | 65.45 | 9.16 |
| GLS (%) | -19.07 | 2.66 | -19.65 | 4.71 | -17.86 | 5.06 | -19.78 | 3.70 |
| GCS (%) | -21.45 | 3.73 | -21.47 | 2.15 | -19.61 | 3.25 | -22.01 | 5.01 |
| | **STRN3$^{+/+}$ /AcPBS (n=13)** | | **STRN3$^{+/-}$ /AcPBS (n=9)** | | **STRN3$^{+/+}$/AngII (n=14)** | | **STRN3$^{+/-}$/AngII (n=10)** | |
| | Mean | SD | Mean | SD | Mean | SD | Mean | SD |
| Heart Rate (bpm) | 517 | 40 | 520 | 46 | 529 | 51 | 517 | 31 |
| Stroke volume (μl) | 28.78 | 5.20 | 28.15 | 5.31 | 27.87 | 4.67 | 25.41 | 5.72 |
| Cardiac output (ml/min) | 14.85 | 2.72 | 14.80 | 4.11 | 14.69 | 2.59 | 13.08 | 2.75 |
| Ejection fraction (%) | 58.04 | 6.77 | 61.92 | 3.13 | 58.94 | 7.01 | 62.44 | 6.16 |
| Fractional shortening (%) | 31.61 | 7.49 | 34.72 | 4.31 | 31.84 | 7.56 | 33.24 | 7.33 |
| EDV (μl) | 50.60 | 13.75 | 45.20 | 7.87 | 47.45 | 8.97 | 40.79 | 9.88 |
| ESV (μl) | 21.82 | 9.25 | 17.05 | 3.08 | 19.57 | 6.22 | 15.38 | 5.34 |
| EDLVM (mg) | 53.99 | 6.23 | 53.83 | 5.42 | 68.06 | 7.48 | 69.43 | 7.30 |
| ESLVM (mg) | 57.12 | 7.14 | 56.01 | 5.81 | 71.12 | 7.46 | 73.15 | 7.28 |
| GLS (%) | -19.46 | 1.96 | -20.56 | 2.92 | -18.79 | 4.11 | -22.72 | 4.00 |
| GCS (%) | -20.39 | 3.51 | -23.53 | 3.81 | -22.95 | 2.46 | -22.80 | 3.68 |

**Supplementary Table S6. Echocardiography data for STRN3$^{fl/fl}$/Cre$^{+/-}$ mice.** Male mice (8 wks) were treated with corn-oil vehicle (COV) or 40 mg/kg tamoxifen (Tx) 4 days before minipumps were implanted for delivery of acidified PBS (AcPBS) or 0.8 mg/kg/d AngII in AcPBS for 7 d. Long axis B-mode images were analysed using speckle-tracking software. Tx, tamoxifen; EDV, End diastolic volume; ESV, End systolic volume; EDLVM, End diastolic left ventricle mass; ESLVM, End systolic left ventricle mass; GLS, global longitudinal strain; GCS, Global circumferential strain.

| | **COV/AcPBS** | | **Tx/AcPBS** | | **COV/AngII** | | **Tx/AngII** | |
|---|---|---|---|---|---|---|---|---|
| | Mean | SD | Mean | SD | Mean | SD | Mean | SD |
| **Baseline** | | | | | | | | |
| Heart Rate (bpm) | 460 | 27 | 465 | 25 | 468 | 32 | 476 | 26 |
| Stroke volume (µl) | 33.26 | 5.25 | 34.35 | 4.56 | 32.10 | 2.68 | 32.60 | 4.42 |
| Cardiac output (ml/min) | 15.36 | 2.82 | 15.96 | 2.23 | 15.04 | 1.89 | 15.43 | 1.67 |
| Ejection fraction (%) | 56.56 | 6.96 | 56.64 | 3.44 | 57.21 | 4.71 | 58.42 | 5.15 |
| Fractional shortening (%) | 30.20 | 6.91 | 30.39 | 3.74 | 31.37 | 4.54 | 31.26 | 4.14 |
| EDV (µl) | 58.76 | 6.92 | 60.69 | 8.28 | 56.20 | 4.25 | 56.26 | 9.59 |
| ESV (µl) | 25.50 | 5.09 | 26.34 | 4.57 | 24.11 | 3.79 | 23.65 | 6.47 |
| EDLVM (mg) | 54.62 | 3.78 | 53.42 | 3.39 | 52.82 | 2.94 | 52.05 | 3.08 |
| ESLVM (mg) | 57.36 | 3.85 | 56.82 | 4.77 | 54.63 | 3.53 | 54.95 | 3.08 |
| GLS (%) | -19.96 | 2.84 | -20.84 | 2.71 | -19.27 | 2.64 | -20.92 | 3.20 |
| GCS (%) | -21.19 | 3.27 | -21.68 | 2.51 | -21.65 | 2.71 | -21.15 | 2.96 |
| **7 d** | | | | | | | | |
| Heart Rate (bpm) | 514 | 56 | 496 | 38 | 490 | 59 | 515 | 62 |
| Stroke volume (µl) | 33.40 | 6.21 | 33.09 | 4.79 | 27.74 | 7.67 | 28.14 | 5.35 |
| Cardiac output (ml/min) | 17.26 | 3.90 | 16.34 | 2.16 | 13.45 | 3.74 | 14.30 | 2.12 |
| Ejection fraction (%) | 60.64 | 8.77 | 59.54 | 5.69 | 56.44 | 11.08 | 59.95 | 6.41 |
| Fractional shortening (%) | 33.88 | 5.48 | 31.30 | 5.47 | 31.45 | 8.26 | 36.14 | 6.01 |
| EDV (µl) | 54.94 | 8.31 | 55.63 | 9.49 | 50.47 | 13.99 | 47.69 | 12.79 |
| ESV (µl) | 21.54 | 6.55 | 22.54 | 6.21 | 22.73 | 8.55 | 19.55 | 7.94 |
| EDLVM (mg) | 55.58 | 4.65 | 54.17 | 4.56 | 71.19 | 15.02 | 58.98 | 4.37 |
| ESLVM (mg) | 58.04 | 5.17 | 57.22 | 4.79 | 72.26 | 15.07 | 60.15 | 4.38 |
| GLS (%) | -20.57 | 6.10 | -20.37 | 3.09 | -16.90 | 3.76 | -16.74 | 3.10 |
| GCS (%) | -23.04 | 4.23 | -22.75 | 3.28 | -22.03 | 3.48 | -21.28 | 5.52 |

**Supplementary Figure S1. Full images for immunoblots in Figure 1.** Proteins were separated on 10% polyacrylamide gels. Red boxes highlight the bands of interest.

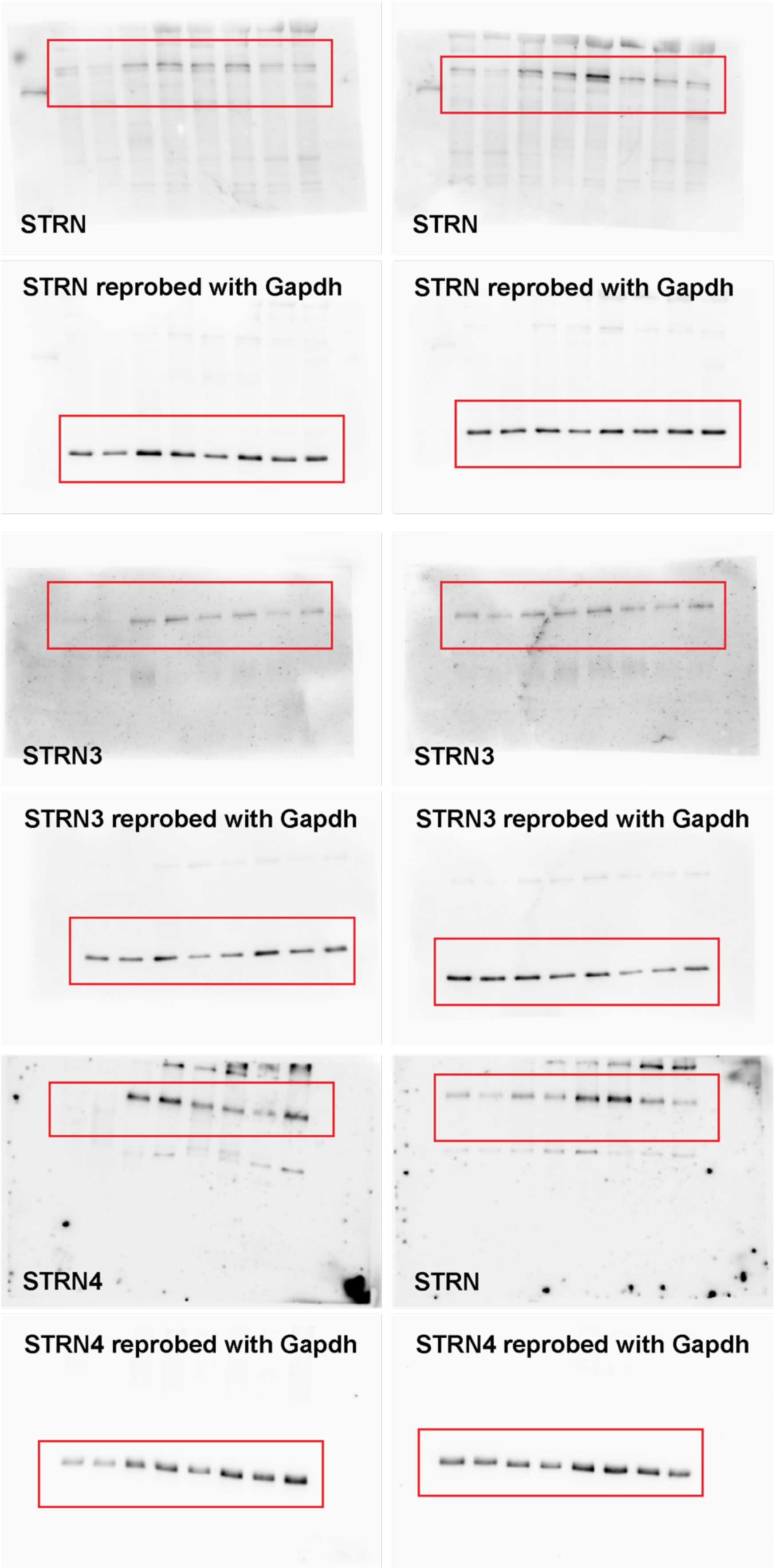

**Supplementary Figure S2. Full images for immunoblots in Figure 2.** Proteins were separated on 8% and 12% polyacrylamide gels for striatins and GAPDH, respectively. Red boxes highlight the bands of interest.

Blots from 2A:

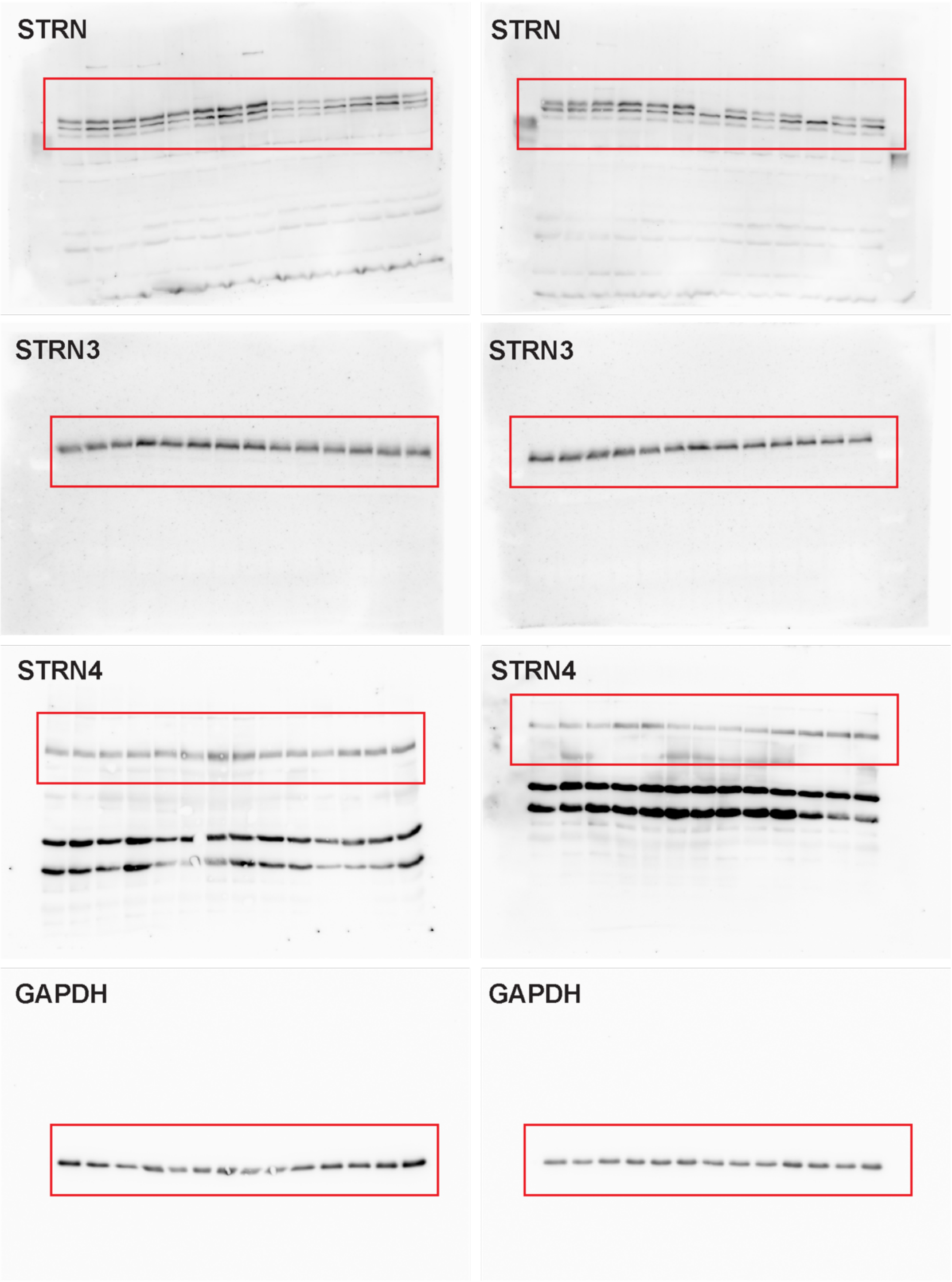

Blots from 2C:

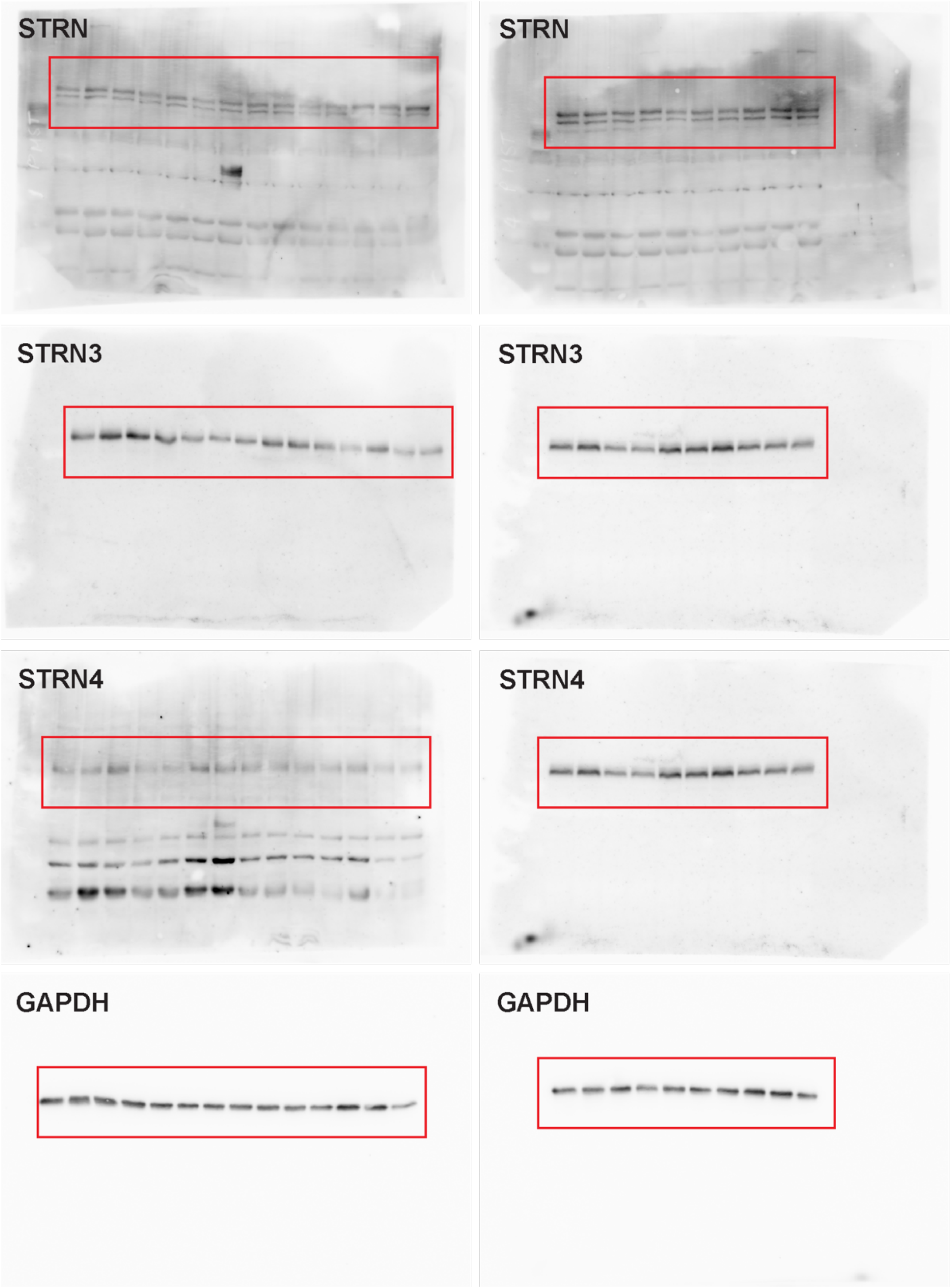

**Supplementary Figure S3. Full images for immunoblots in Figure 5.** Proteins were separated on 8% and 10% polyacrylamide gels for striatins and GAPDH, respectively. (N.B. Two half blots for GAPDH from different experiments were imaged together giving upper and lower bands). Red boxes highlight the bands of interest.

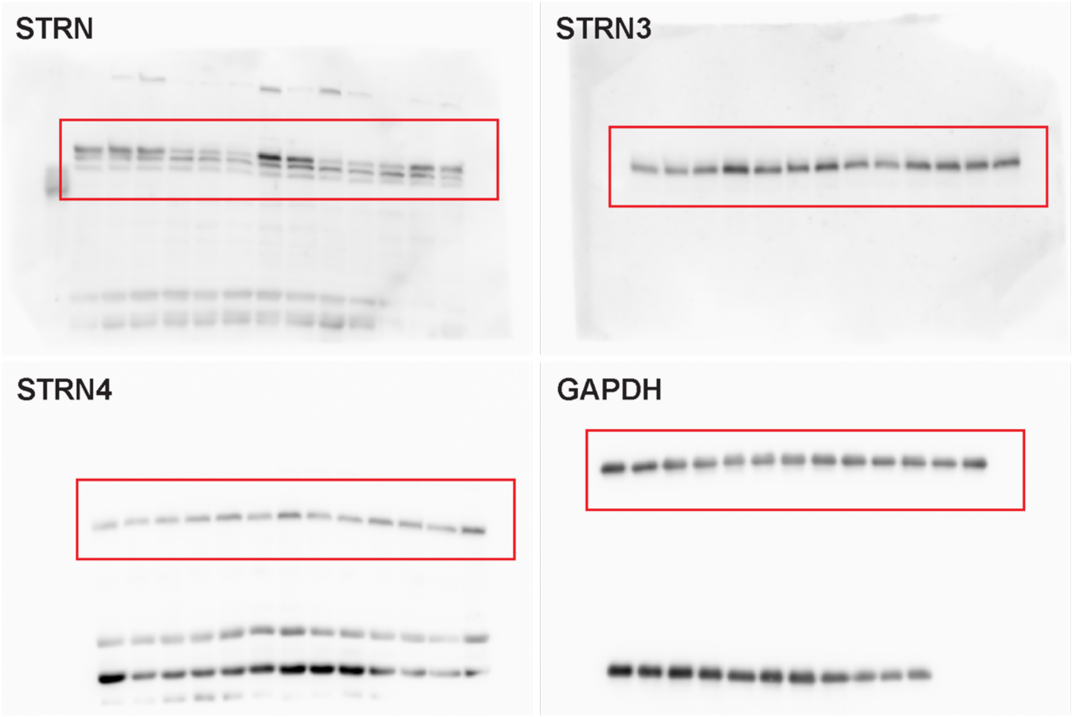
